# FRET-sensitized acceptor emission localization (FRETsael) – nanometer localization of biomolecular interactions using fluorescence lifetime imaging

**DOI:** 10.1101/2023.12.10.570984

**Authors:** Yair Razvag, Paz Drori, Shalhevet Klemfner, Eran Meshorer, Eitan Lerner

## Abstract

Super-resolution light microscopy techniques facilitate the observation of nanometer-size biomolecules, which are 1-2 orders of magnitude smaller than the diffraction limit of light. Using super-resolution microscopy techniques, it is possible to observe fluorescence from two biomolecules in close proximity, however not necessarily in direct interaction. Using FRET-sensitized acceptor emission localization (FRETsael), we localize biomolecular interactions exhibiting FRET with nanometer accuracy, from two-color fluorescence lifetime imaging data. The concepts of FRETsael were tested first against simulations, in which the recovered localization accuracy is 20-30 nm for true-positive detections of FRET pairs. Further analyses of the simulation results report the conditions in which true-positive rates are maximal. We then show the capabilities of FRETsael on simulated samples of Actin-Vinculin and ER-ribosomes interactions, as well as on experimental samples of actin-myosin two-color confocal imaging. Conclusively, the FRETsael approach paves the way towards studying biomolecular interactions with improved spatial resolution from laser scanning confocal two-color fluorescence lifetime imaging.

**Significance:** FRET is used in fluorescence microscopy to report whether dye-labeled biomolecules of choice are close within distances of 10 nm or less, hence typical interaction distances. However, in many cases, using FRET imaging for the study of biomolecular interactions is difficult due to the high density of dye-labeled biomolecules and due to the existence of unbound dye-labeled biomolecules. In addition, the resolution of localizing molecules using light microscopy is diffraction limited. This work presents *FRETsael*, a new approach for localizing interacting biomolecules undergoing FRET, with improved resolution of 20-30 nm for confocal microscopy using search algorithms for local extrema in contribution to FRET using two-channel fluorescence intensity and lifetime data.

## Introduction

Since its conception, super-resolution (SR) light microscopy has aimed at observing biomolecules in cells, and by that achieve the maximal information content in space and strive to achieve them in real time (1, 2). The sizes of biomolecules that drive the processes of life are as small as a few nanometers (nm) for small proteins with a few domains, and as large as tens of nm in width and hundreds of nm in length for polymeric ultra-structures such as the chromatin in the cell nucleus, cytoskeletal fibrils in the cytoplasm or protein aggregates and perhaps even amyloid-like fibrils at different cellular locations. Viewing biomolecules in live cells can be performed using light microscopy, which is diffraction-limited. Hence, when light in the visible range is used, it is limited to spatial resolutions that are 1-2 orders of magnitude larger relative to biomolecules’ sizes. Indeed, using electron microscopy solves this problem and allows recovering the fine biomolecular details of fixated or cryo-frozen cells, but does not allow observing these details in live cells or in ambient conditions. Therefore, studying biomolecular interactions in the cell at nanoscale resolution, while biological activity is occurring, is difficult using light microscopy.

SR microscopy assists viewing the nm-scaled biomolecular world of the cell, focusing on a subset of biomolecules that were labeled with fluorescent proteins (FPs), dye-labeled antibodies, dye-labeled oligonucleotide probes or clickable organic dyes. The diffraction limit of light is a physical boundary, and hence cannot be directly overcome. However, SR microscopy techniques provide ways to gain the nm spatial resolution by introducing additional molecular contrast on the imaged items by using special features of the fluorescent dyes, such as on/off fluorescence blinking, which is at the heart of single molecule localization microscopy (SMLM) techniques (3–5), or by using special optics, such as in stimulated emission depletion (STED) microscopy (6–10) or minimal flux (MINFLUX) microscopy (11–13).

Imaging interactions between different biomolecules is more complex. Dual-color fluorescence imaging of two FPs is a very common method in cellular imaging, and is typically used for reporting the localizations of these interactions with accuracies of >300 nm (14). However, such co-localizations do not necessarily report the nm proximities required for maintaining direct biomolecular interactions between biomolecules.

Another possibility is to utilize Förster resonance energy transfer (FRET) (15–17). In FRET, the excitation energy of a donor dye can be transferred to an acceptor dye with a given efficiency that inversely depends on the donor-acceptor distance. The donor-acceptor distances accessible by FRET are <10 nm, which is a typical range for the distance between two FPs fused to two proteins that are in direct contact due to an interaction (18). By combining FRET with SR methods, biomolecular interactions were resolvable with excellent localization accuracies and precisions (19–22). It would be beneficial, however, to attain localizations of biomolecular interactions in a simple confocal microscope setup and with common fluorescent dyes or tags by using the knowledge of the non-uniform Gaussian-shaped approximate illumination profile.

To achieve this goal, we do not only use fluorescence intensity variations due to FRET but utilize another dataset that is achievable when using a confocal based system, fluorescence lifetimes of the donor or acceptor dyes. In this study, the molecular-level contrast, i.e. the sparsity of molecules to achieve SR, is plausible between direct interactions of proteins with fluorescent tags (e.g., fused to FPs, fluorescently-labeled antibodies) that are within FRET proximities, and the same proteins that are not interacting and hence are not within FRET proximities.

The common practice in using confocal imaging is to laser-scan a given X-Y plane in the sample, with scan steps on the scale of half the excitation wavelength. We perform fluorescence lifetime imaging (FLIM) (23–26) and laser-scan images using smaller (e.g., few nm) step sizes, and then localize the pixels in which the contribution to FRET, as was assessed by several fluorescence-based parameters, was locally maximal. Therefore, we termed the method FRET-sensitized acceptor emission localization, or *FRETsael*. We quantify the contribution to FRET based on several fluorescence-based parameters, such as the acceptor fluorescence lifetime after donor excitation, relative to the acceptor fluorescence lifetime after acceptor excitation, in a pulsed-interleaved excitation (PIE) (27) layout (also known as nanosecond alternating laser excitation, nsALEX layout (28)), which reports the delay in the acceptor fluorescence decay due to FRET-sensitized acceptor emission. To test the FRETsael approach and characterize its strengths and limitations, we perform a battery of tests against simulated data with known ground-truth, in which we define the localization accuracies and sensitivities of FRET localizations. Then, we demonstrated the capabilities of FRETsael on simulations based on previously-acquired electron microscopy images of (i) actin and vinculin and (ii) endoplasmic reticulum (ER) and ribosomes, for which we introduced localizations of ER-ribosome interactions as well as of non-interacting ribosomes, depicting the extra information the method can provide. Finally, we took the FRETsael approach into the cell, where ground-truth for each dye-labeled biomolecule is not known, to probe the well-defined interaction between actin and myosin.

## Results

### The FRETsael concept

The reader is referred to the supporting text for an in-depth expansion of the presented approach and system. While the herein presented approach is relevant for FRET imaging of any dye-labeled biomolecule, for the sake of simplicity the definitions here will focus on proteins as the dye-labeled biomolecules. However, the approach can be used also with dye-labeled DNA, RNA and other biomolecules suspected of undergoing direct interactions. One can envision the imaged part of the sample, using a given scanning step size in a laser scanning confocal microscope (e.g., Fig. 1, A), as including (i) a variety of proteins labeled with a donor dye of type “P_D_”, and (ii) other proteins labeled with an acceptor dye of type “P_A_”, where (iii) a fraction of these proteins associate to form an interacting complex P_D-A_, and that (iv) the D-A distance is within the range in which FRET occurs, while (v) the distance between these proteins, when they do not interact, is typically larger than the FRET range (see Fig. 1, B, as a possible layout).

**Fig. 1.**
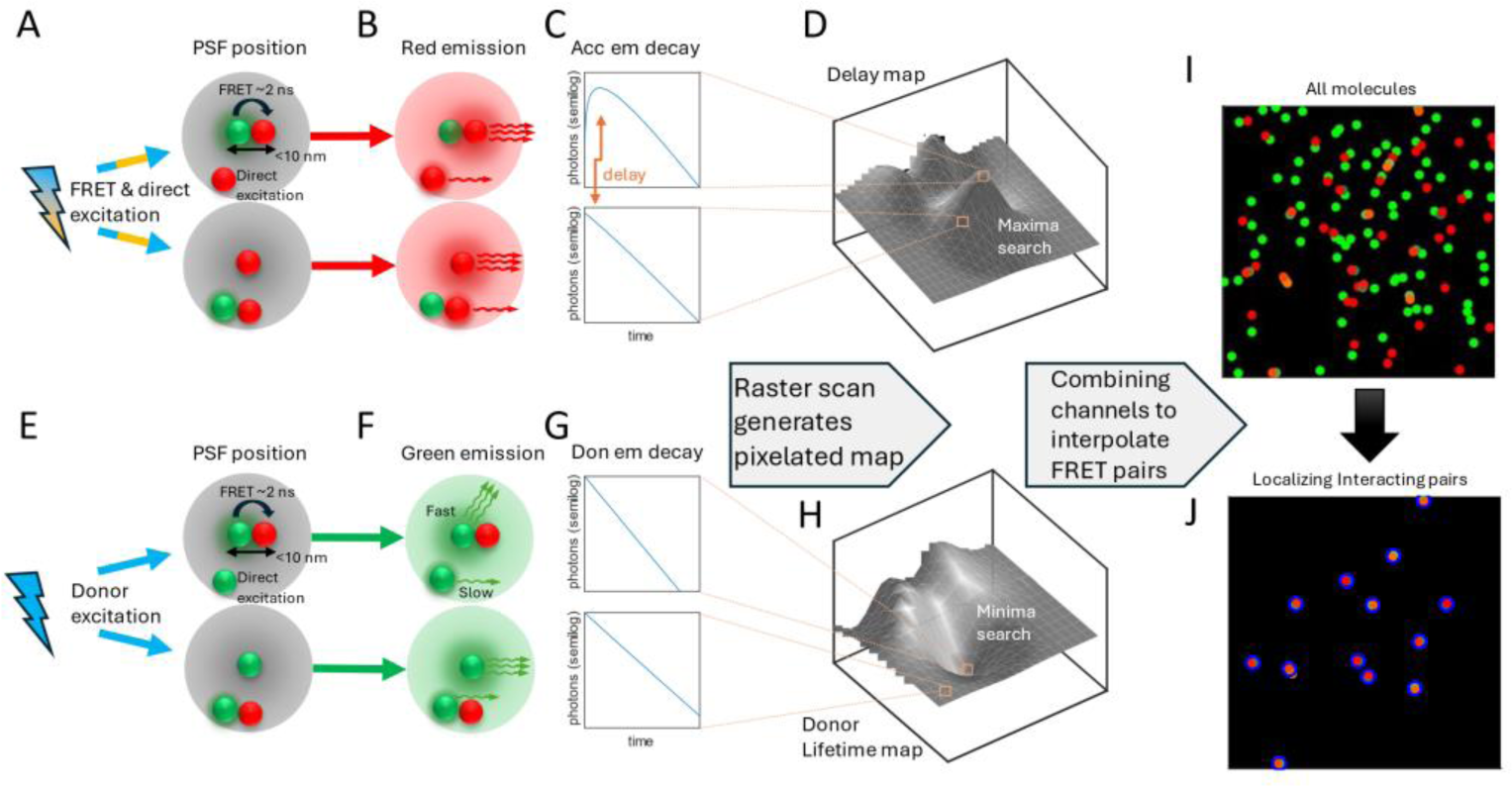
The concept of FRETsael using laser scanning confocal-based PIE-FLIM setup. Illustration demonstrating two of the four parameter channels used for localizing interacting molecule pairs in FRETsael; **A**. Description of parameter channel 1: delay in acceptor mean fluorescence lifetime due to contribution from FRET. Confocal based system enables raster scanning the sample, thus at some point, interacting pair will localize at the center of the PSF (upper panel; out of scale) or at the periphery of the PSF (bottom panel). Acceptor (red) excitation occurs either via energy transfer by exciting the donor (green) fluorophore first (cyan laser) or via direct excitation (orange laser). **B.** Acceptor photons (red) are emitted at specific times to generate an acceptor fluorescence decay plot depicted in **C**, whereas as interacting pair located in the center of the PSF will dominate photon emission, thus the decay will have a delay in time due to the time spent in the donor before FRET occurred. **D.** Raster scanning the sample by using small laser scanning steps will provide a delay map by subtracting the average acceptor lifetime after direct excitation from the average acceptor lifetime after donor excitation. **E.** Description of parameter channel 2: donor fluorescence lifetime reduction. In this parameter channel we separate cases where the interacting pair (upper panel) or a donor-only molecule (bottom panel) are centered at the center of the PSF. **F.** Donor molecule (green) is excited (cyan laser) and will emit donor fluorescence photons (green). In parallel if there is an acceptor in FRET proximity, it can also transfer the excitation energy via FRET, which will lead to a reduction in the donor fluorescence lifetime. **G.** Donor fluorescence decay plots. In the case of an interacting pair located at the center of the PSF, the decay will be faster as the longer time the fluorophore will stay in the excited state, the higher the probability that the energy will be transferred to the acceptor (upper versus bottom panels). **H.** Raster scan with small laser scanning steps provides a donor fluorescence decay lifetime map. **I.** and **J.** By combining information from all parameter channels, i.e. local maxima search in parameter channel 1 (**D**) or local minima search in parameter channel 2 (**H**), a localization map is created (**J**) which provide localizations of interacting pairs at improved resolution, out of the dense sample of all donors and acceptor molecules (**I**).

In this illustration, each protein or protein complex occupies a given self-size, of a few nm, typical of most proteins. Assuming the concentration of these proteins at the imaged region is high, but not to the level in which proteins P_D_ and P_A_ will occasionally exist in <10 nm distances for prolonged periods of time, we can assume that most contributions to FRET would arise from complexes P_D-A_. Additionally, we assume that the proteins P_D_ and P_A_ at the imaged region are immobile at least at the timescales of the acquisition of that pixel, which would be the case for relatively slow moving proteins or alternatively with fixated samples. The excitation profile can then be added, which can be approximated by a Gaussian shape. To an approximation, the value of each point along the Gaussian-shaped point-spread function (PSF) can be conceived as the photon flux for excitation at this point. With this description at hand, we could envision the following scenarios:

1. The maximum of the Gaussian-shaped PSF is positioned right above a donor dye attached to protein P_D,_ in the complex P_D-A_, and there are other proteins P_D_ and P_A_ that are far apart and do not contribute to FRET, found at positions on the periphery of the Gaussian-shaped PSF. In this case, there will be different contributions to donor excitation, where (i) the ones that do not contribute to FRET will emit donor photons detected in the donor channel, (ii) perhaps a small fraction of that non-FRET contribution will leak into the acceptor detection channel, and (iii) the predominant contribution to FRET, and hence to the emission in the acceptor detection channel, would arise from the PSF-centered P_D-A_ complex.
2. The maximum of the Gaussian-shaped PSF has now moved a few nm away from the P_D-A_ complex, and so the excitation rate of its donor will decrease, which will reduce the amount of contributions to FRET on the expense of a potential increase in the contributions to excitation of other donor-labeled P_D_ molecules that do not exhibit FRET, due to large distances from non-interacting acceptor-labeled P_A_ molecule.

This concept can also be shown in 1D, for the sake of simplicity (Fig. 2).

**Fig. 2.**
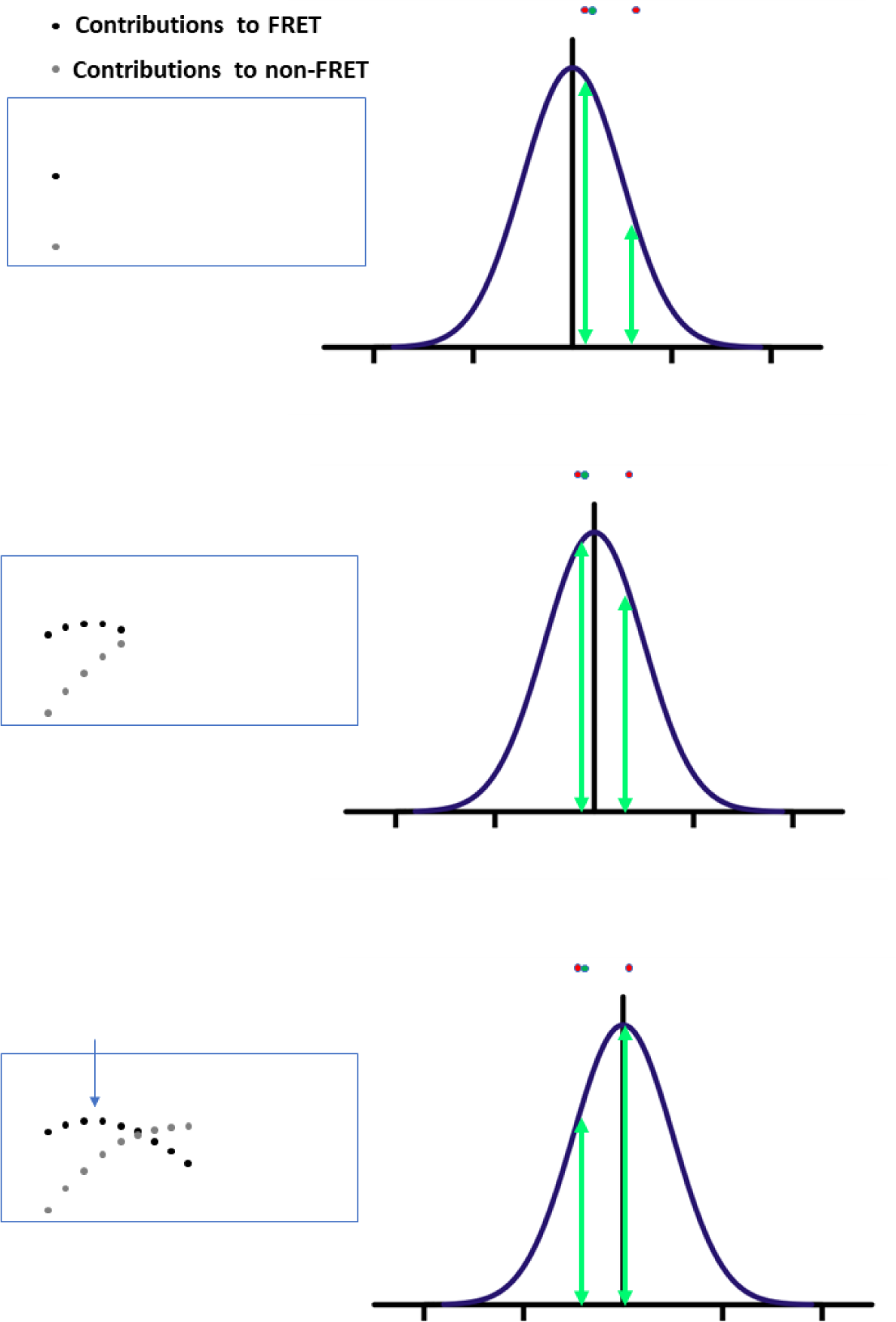
The concept of FRETsael in 1D. A 1D Gaussian profile is scanning two biomolecular entities found at different locations in small steps, one is not involved in FRET (red sphere) and one is involved in FRET (green and red interacting spheres), and the contribution to their excitation is illustrated in each scanning step as vertical green arrows. The contributions to FRET and to non-FRET are presented in the insets in black and grey dots, respectively, for each scanning step. All that is left is to seek the local maximum of the contributions to FRET, using a proper parameter that reports on FRET unambiguously.

What is left is to precisely quantify the contribution to FRET in each such pixel and to spatially identify the local maxima of such contributions. One can record the donor and acceptor fluorescence intensities using two point-detectors optically filtered for donor and acceptor fluorescence. However, this approach assumes that all changes in donor or acceptor fluorescence intensities are solely due to changes in FRET. There are other sources of variability in donor and acceptor fluorescence intensities that are not related to FRET, such as variation in the fluorescence quantum yields of the donors or acceptors dyes at different intracellular environments with different physico-chemical conditions.

Another approach is to employ fluorescence lifetime imaging (FLIM) capabilities and then report reductions in donor fluorescence lifetimes, τ_D_, as reporters of FRET (see eq. S21 in supplementary text). However, there are reasons other than FRET for variability of donor fluorescence lifetimes, and therefore with this approach there might be no stable reference value for donor fluorescence lifetimes. A common approach to tackle this issue is to first record the donor fluorescence lifetime data in the presence of the acceptor, τ_DD_, then repeat this acquisition following acceptor photobleaching, to recover the reduction of the donor fluorescence lifetimes relative to the fluorescence lifetime values in the absence of photo-active acceptor dyes, τ_D_, in each pixel (29, 30) (see eq. S21 in supplementary text). However, this approach is irreversible per the imaged region, and oftentimes does not provide the required full acceptor photobleaching or lack of donor photobleaching for that matter. One can report the acceptor mean fluorescence lifetime after donor excitation, τ_DA_, which should depend on the time in which the excitation energy stays in the donor excited-state, and then in the acceptor excited-state, in the well-defined process known as FRET-sensitized acceptor emission (31–34) (see eq. S22 in supplementary text). This experimental parameter could have served as an excellent reporter of FRET (i) if only the excitation source that leads to donor excitation would not have induced also a fraction of acceptor excitation, and (ii) if only the donor fluorescence would have not leaked into the acceptor fluorescence detection channel (these non-FRET contributions are explained in the supplementary text and in eqs. S14, S15). In practice, in most organic dyes and FPs, it is difficult to find an available laser wavelength for excitation that will solely excite the donor dye. Additionally, the long tail of fluorescence spectra of most organic dyes and FPs almost always leaks to the acceptor fluorescence detection channel.

From the imaging standpoint, a donor excitation source that excites an acceptor molecule directly or donor fluorescence leakage into the acceptor detection channel, per a single acceptor or donor molecule, do not contribute much relative to the photons arising from FRET-sensitized acceptor emission. However, if many such donors or acceptors are found in the region of excitation that include also the donor-acceptor FRET pair, P_D-A_, they will contribute many FRET-irrelevant acceptor photons. A situation, in which one FRET pair is imaged in the presence of thousands of donor-or acceptor-labeled molecules not involved in FRET, is certainly possible in FRET cellular imaging. Importantly, however, the main difference between the FRET-sensitized acceptor emission photons and the non-FRET photons can be identified in fluorescence lifetimes, where the former FRET process has a longer and delayed acceptor fluorescence decay (see eq. S14) relative to the latter non-FRET processes (see eqs. S17, S18). Additionally, donor-only and acceptor-only molecules, P_D_ and P_A_, respectively, will add non-FRET signals. Therefore, each pixel in the image will include donor or acceptor fluorescence lifetime information that are a superposition of the FRET-relevant and FRET-irrelevant (i.e., FRET and non-FRET) processes. Additionally, this superposition of fluorescence signals is weighted by the positions of the molecules relative to the excitation profile (eqs. S19, S20). Relying on the spatially non-uniform Gaussian-shaped excitation profile, and the self-volume of each such protein, there will be a position relative to the excitation PSF, in which the FRET pair in the protein complex P_D-A_ will be aligned with the maximal position of the Gaussian, and there, the delay in the overall acceptor fluorescence decay will be maximal, relative to this value at nearby pixels (Figs. 1, 2).

Following the above concepts, the experimental contribution to FRET per each pixel will be quantified based on mean fluorescence lifetime and fluorescence intensity ratio parameters (eqs. S21-S25, S28), and then local extrema of these parameters will be identified to report local maxima of contributions to FRET. Doing so, the accuracy of localizing FRET from a P_D-A_ complex will depend on the spatial contrast between a density of P_D-A_ complexes and a density of P_D_ and P_A_ proteins, as well as the size of the laser scanning step – the smaller the step is, the finer the changes of the Gaussian excitation profile relative to the positions of P_D-A_ complexes and P_D_ and P_A_ molecules are.

Importantly, however, much like it is for donor fluorescence lifetimes, there are also reasons for variability in acceptor fluorescence lifetimes that are irrelevant to FRET. Therefore, it is important to compare the pixel-based *τ_DA_* to the actual acceptor fluorescence lifetime after direct acceptor excitation, *τ_AA_*, as rapidly as possible. We do so by employing the PIE configuration, in which two pulsed laser excitations, one with a wavelength optimal for donor excitation, and another with a wavelength optimal for acceptor excitation, are interleaved so that the fluorescence decay following donor excitation (including the FRET-sensitized acceptor emission) will be followed by the acceptor pulsed excitation and the fluorescence decay following acceptor excitation. This way, each pixel that is imaged for a few milliseconds (ms), undergoes multiple donor and acceptor interleaved excitations, leading to multiple detection times of (i) donor photons after donor excitation, *D_ex._D_em._*, (ii) acceptor photons after donor excitation, *D_ex._A_em._* and (iii) acceptor photons after acceptor excitation, *A_ex._A_em._* (35). Then, the *A_ex._A_em._* fluorescence lifetime, *τ_AA_*, can be subtracted from the *D_ex._A_em._* fluorescence lifetime, *τ_DA_*, to gain a relative measure of the contribution to FRET within a pixel (see also eq. S24 and the supplementary text). Thus, our main parameter channel of interest is (eq. 1)

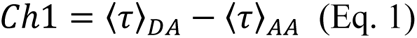

which describes the delay in acceptor mean fluorescence lifetime due to the contribution from FRET sensitization of acceptor fluorescence (see full derivation in eqs. S22-S24). Three other parameter channels we chose to focus on, which are indicative of contributions to FRET, are the donor fluorescence intensity and fluorescence lifetime, both decreasing, due to FRET (eqs. 2, 3; notably, the numbering of the parameter channels is according to how they are presented in the results section; see full derivation in eqs. S21, S25)

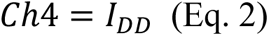

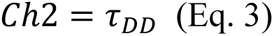

and the ratio of acceptor and donor fluorescence intensities after donor excitation (eq. 4; see full derivation in eqs. S26, S28).

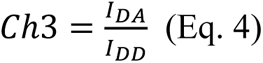

Finally, an image of these parameter channels is blurred, due to the use of small laser-scan steps, because in each laser scanned pixel, the excitation beam differently excites all the fluorophores coinciding within the Gaussian-shaped PSF. The local extremum values in the image depend on the parameter channel, which will guide the inference of the localization of maximal contributions to FRET. Thus, as a final step, a local extremum search algorithm is applied to images of the parameter channels.

### Testing FRETsael in simulations against ground-truth

To test the accuracy of localizing FRET contributors as well as the sensitivity of detecting them, we first examine the algorithm *in silico* through a set of simulations. Doing so, we could test the implementation of the FRETsael concept against known ground-truth localizations of donor– and acceptor-labeled proteins undergoing FRET. Furthermore, we can test the performance of FRETsael under various conditions.

Therefore, using Monte Carlo (MC) calculations we simulated the laser scanning confocal imaging of locations of donors and acceptors in an X-Y plane (Fig. S1), under different conditions. For the simulation of FRET imaging, we used the literature parameters of eGFP and mCherry as a common FP-based FRET pair (36, 37).

Then, since the ground-truth locations of all donors and acceptors are known, we calculated the localization accuracy for all true positive (TP) events (i.e., where ground-truth P_D-A_ FRET pairs was identified) as the distance between the ground-truth position of the P_D-A_ and the local extremum-based localization. Additionally, we identified false positive (FP) events as local extrema localizations, in which no ground-truth FRET pairs existed, and false negative (FN) events as ground-truth FRET pairs that were not found as local extrema (Fig. S2).

Using these TP, FP and FN events, we calculated the false discovery and true positive rates (FDR and TPR, respectively) per each simulation condition tested (eqs. 5, 6).

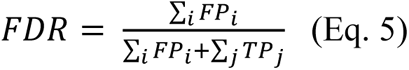

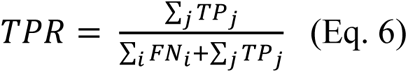

where the ideal result is reached if the FDR reaches a value of zero and the TPR reaches a value of one. To combine the three event types, TP, FP and FN, into one function, the critical success index (CSI) (38) was used (eq. 7).

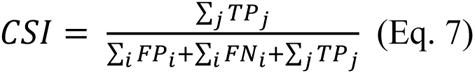

where a value of one represents the perfect result and zero the poorest.

As described above, we studied four different FRET reporter parameter channels (see eqs. 1-4). Then, we searched these images to detect local maxima in parameter channels 1 and 3, or local minima in parameter channels 2 and 4 (Fig. S1). The results of the simulations for varying (a) donor-acceptor stoichiometries, (b) donor and acceptor densities, (c) strength of interaction, (d) widths of the excitation Gaussian profile and (e) laser scan step sizes are summarized in Fig. 3.

**Fig. 3.**
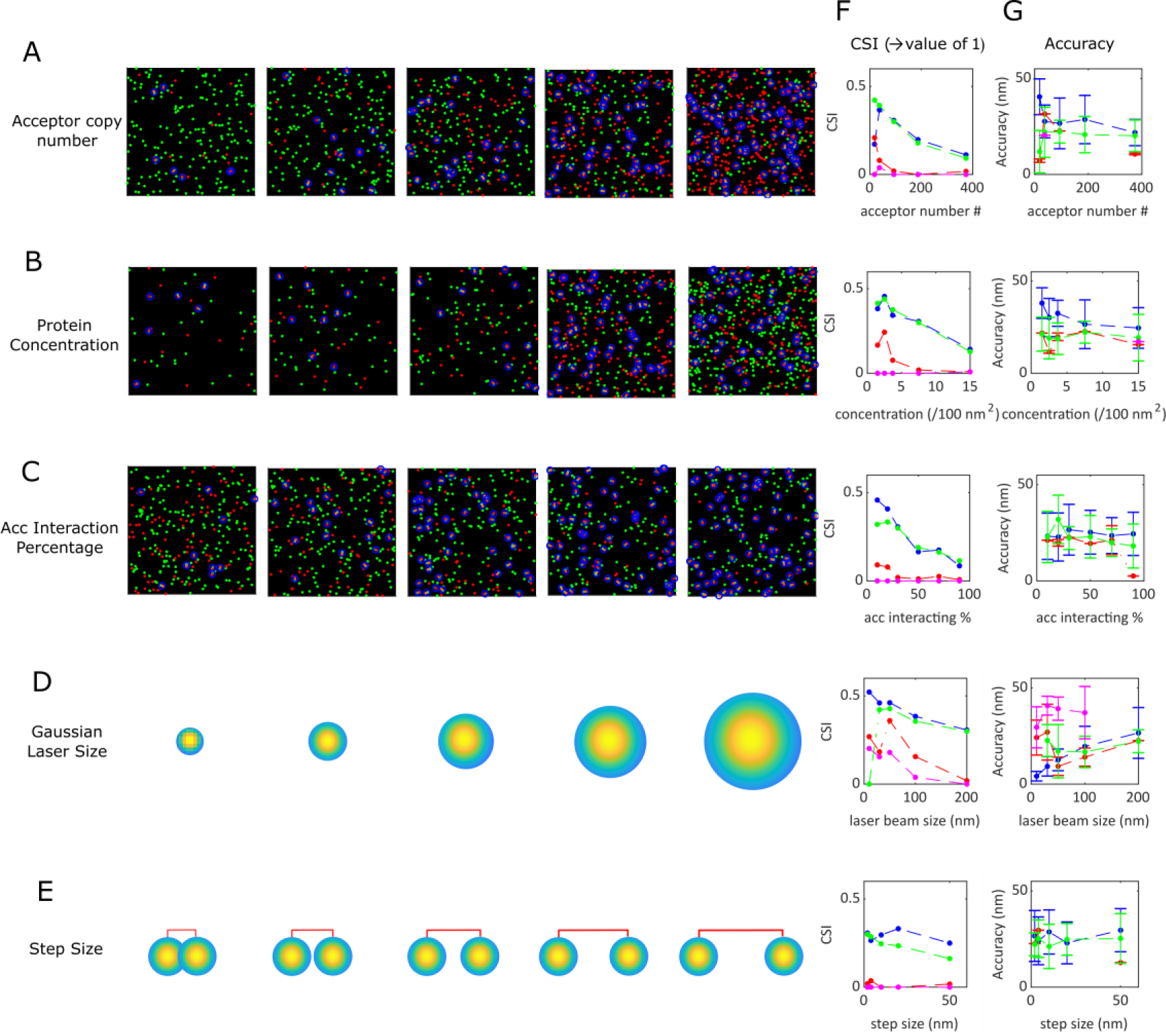
Testing FRETsael on simulated data against known ground-truth. **A-E**. Simulated ground-truth positions of donor (in green), acceptor (in red) and FRET pairs (mark in purple), in a 500×500 nm^2^ frame, with varying conditions for the different simulated conditions: donor-acceptor stoichiometries (**A**), donor and acceptor densities (**B**), strength of interaction (**C**), widths of the excitation Gaussian profile (depiction) (**D**), and laser scan step sizes (**E**; depiction). **F**. The values of the critical success index (CSI) for the different condition values tested in panels **A**-**E**. Each channel is depicted with a different color; the delay in the acceptor mean fluorescence lifetime due to contribution from FRET (blue), the donor mean fluorescence lifetime (red), the ratio of acceptor and donor fluorescence intensities after donor excitation (green) and the donor fluorescence intensity (pink), as well as the mean between all four parameters (black). **G**. The accuracy of FRET localizations in TP events for local extrema identified based on images for the four different channels.

Interestingly, using a Gaussian-shaped PSF with widths like those achieved in confocal imaging, the localization accuracy was 20-30 nm for all TP events, regardless of the simulation conditions (Fig. 3). In fact, reducing the width of the Gaussian-shaped PSF to 50 nm, which is achievable in STED microscopy, led to even improved localization accuracies of <20 nm (Fig. 3, D, G). However, the sensitivity to detect FRET localizations is also important. In that respect, the densities or concentrations of the donor and acceptor molecules, as well as their stoichiometries should play a key role in the sensitivity. We found that the CSI decreases as the donor:acceptor stoichiometry decreases. However, a donor:acceptor stoichiometry of two, yielded an optimal decrease in the FDR, while maintaining a good TPR of ∼50% identifications for parameter channels 1 and 3.

In addition, we found that when we increase the total amount of donors and acceptors, but keep their donor:acceptor stoichiometry at a value of two, both the FDR and TPR decrease with increasing overall concentrations, which can also be seen in CSI decreases (Fig. 3, B, F). However, in the range of 1-5 proteins per 100 nm^2^ (equivalent to a few nM concentration), which is a biologically relevant concentration (39), the FDR was kept at ∼30%, while the TPR reached ∼60-70%, at least for some of the parameter channels (the local maximum in the CSI curve; Fig. 3, B, F).

Increasing the interaction strength, as shown by the fraction of donor-acceptor complexes, led to a decrease of CSI values (Fig. 3, C, F), because (a) the concentration of the FRET pairs is increasing, and (b) there are less non-interacting acceptor molecules, which means less molecular contrast between FRET and non-FRET contributions. Within these trends, it was interesting to find that among the different parameter channels, the best performing ones were the delay in acceptor mean fluorescence lifetime due to contribution from FRET (eq. 1), and the ratio of acceptor and donor fluorescence intensities after donor excitation (eq. 3) – lowest FDR values among all parameter channels, and intermediate TPR values among all parameter channels.

Reducing the width of the Gaussian-shaped PSF led to improved localizations with all four parameter channels, as expected, due to the more concise excitation profile. Moreover, using narrower excitation profiles, localization accuracies can be improved to a few nm precision (Fig. 3, D, F, G).

Nevertheless, the condition that was most relevant to FRETsael to be tested was the laser scan step sizes, or in other words, how fine the parameter channel maps should be in order to search local extrema. Interestingly, we found that the laser scan step size is crucial for the acceptor to donor intensity ratio parameter channel (Fig. 3, E, F, green line), since the larger the laser scan step size was, the less TP localizations were identified. As for the parameter channel of the acceptor mean fluorescence lifetime delay an optimal laser scan step size of ∼20 nm was identified (Fig. 3, E, F, blue line).

Noticeably, some parameter channels exhibited improved performance relative to others, i.e., the delay in acceptor mean fluorescence lifetime due to contributions from FRET performed best in most cases, while the donor fluorescence intensity did not at all. It could have been mistakenly assumed that while a FRET pair will result in a long delay in acceptor mean fluorescence lifetime due to contribution from FRET (which can decrease as the distance of the center of the PSF from the FRET pair increases), the high density of donor-only molecules, can reduce this delay. We, however, do not identify this effect in the simulations; the delay caused by the FRET pair is predominant and clear, and reduces upon getting the center of the PSF farther away from the center position of the FRET pair (Fig. S3 A, C, E). On the other hand, in the donor only case, the delay, if any, is unchanging throughout the distance from the position center (Fig. S3 B, D, F).

Nevertheless, although parameter channels 1 and 3 perform much better, the interpolation ability of local extrema can be further improved by gathering information from a combination of parameter channels, which could potentially increase TP events. Therefore, the convention was that a localization is considered only if at least two parameter channels provided a localization in the same area of a circle in a diameter of 50 nm (Fig. 4).

**Fig. 4.**
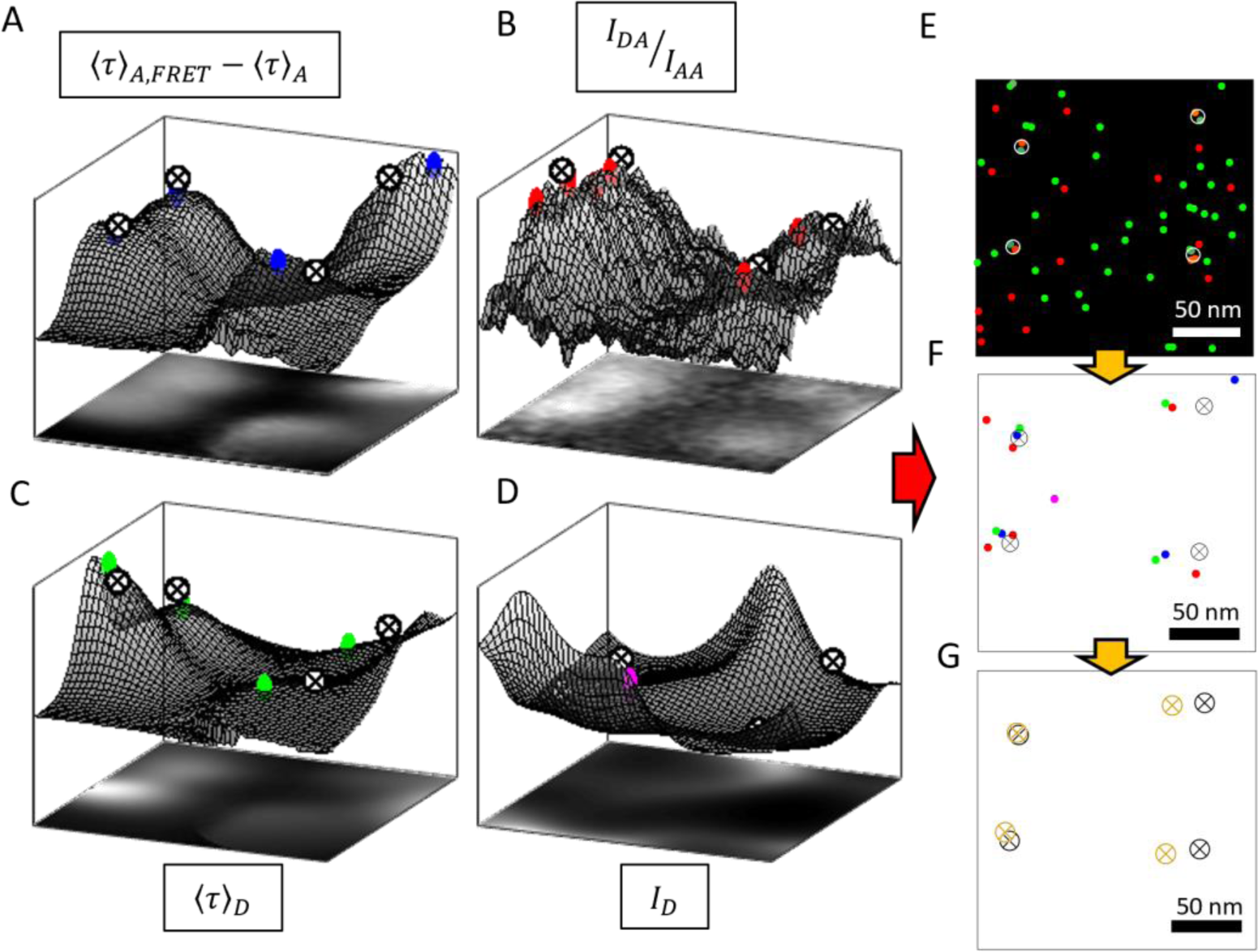
Combination of local extrema localizations of different parameters. **A-D**. Demonstrating the extremum search procedure – topographic maps of all four parameter channels: **A.** the delay in the acceptor mean fluorescence lifetime due to contribution from FRET; **B.** the donor mean fluorescence lifetime; **C.** the ratio of acceptor and donor fluorescence intensities after donor excitation; **D.** the donor fluorescence intensity. **E**. The corresponding simulated molecules for the area depicted in panels **A**-**D**. A total of 40 donor and 20 acceptor molecules were randomly distributed. Among these, four were in FRET proximity, acting as FRET pairs. **F**. Localizations derived from local extrema in maps of the four parameter channels are presented as circles (delay in the acceptor mean fluorescence lifetime in blue; donor mean fluorescence lifetime in red; ratio of acceptor and donor fluorescence intensities after donor excitation in green; donor fluorescence intensity in pink). Ground-truth locations of the four donor-acceptor FRET pairs are presented as black ⊗ signs. **G**. Ground-truth locations alongside localizations after integrated the four different channels (orange ⊗ sign). Simulated 500×500 nm^2^ frame, with a 50 nm scale bar.

While these MC simulations assisted us testing the FRETsael approach, identifying the best parameter channels to use and defining the method limitations, we note that these simulations did not consider the typical inhomogeneous spread of biomolecules inside cellular compartments. This is important to consider, since in many cases different biomolecules tend to localize more in some parts of the cell rather than in others, leading to clear changes in their concentrations, as well as in the concentrations of complexes of these biomolecules with others. In the next section, we test the FRETsael approach on such simulations.

### Testing FRETsael on a simulated ground-truth with a well-defined shape

Next, we sought to simulate a realistic distribution of the donor and acceptor proteins to test the behavior of FRETsael on a sample that could resemble biological scenarios in cellular imaging using simulations. To do so we simulated actin molecules within filaments as donor proteins with vinculin, a protein related to focal adhesion sites, as acceptors (40). To maintain biological relevance, we simulated five filaments over a 1×1 µm^2^ frame, with 150 donor and 40 acceptor molecules, having 60% of them involved in a ground-truth interactions bringing donor-acceptor pairs to FRET distances, thus, in total ∼25 donor-acceptor pairs undergoing FRET due to an interaction. The simulation results show that even for a more realistic layout, the implementation of the FRETsael concept works well and provides a CSI value of 40% with a TPR of ∼58% while keeping the FDR as low as ∼42% with accuracy levels of 51±20 nm (SD; Fig. 5). Importantly, in this simulation the threshold for the parameter channel of the donor intensity was restricted as it introduced elevated levels of noise, as expected because there were many non-interacting donor molecules.

**Fig. 5.**
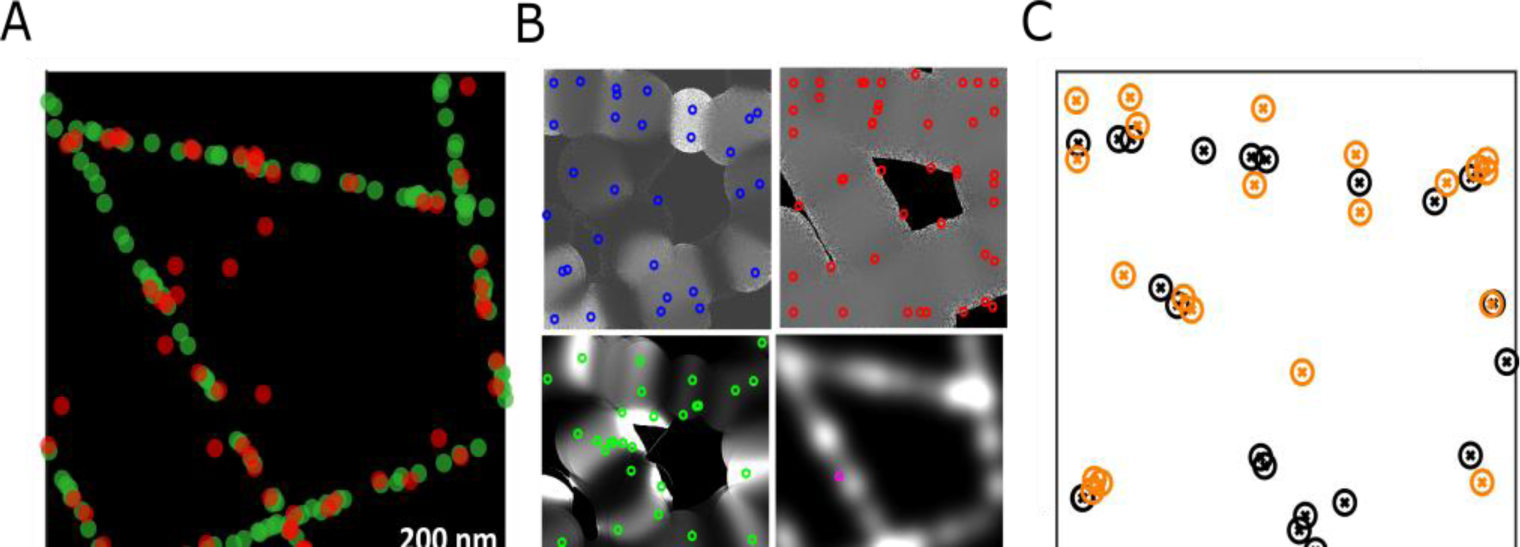
Actin-Vinculin simulation. **A**. Donors and acceptors, representing actin (#150; green dots) and vinculin (#60; red dots), respectively, were distributed over five lines and over a of 1×1 µm^2^ frame. Low percentage of the red vinculin molecules were distributed not over the lines – to introduce non-interacting events, and hence noise into the simulation. **B.** Images of the four parameter channels with localizations of local extrema in each channel (delay in the acceptor mean fluorescence lifetime in blue; donor mean fluorescence lifetime in red; the ratio of acceptor and donor fluorescence intensity after donor excitation in green; donor fluorescence intensity in pink). **C**. The ground-truth of interactions (40% in interactions, leading to 24 FRET pairs) represented as black ⊗ signs. Localizations of suspected interacting pairs (orange ⊗ signs) presented next to the ground-truth locations. Comparing localizations with ground-truth locations yielded detection accuracy values of 51±20 nm (SD) with TPR of 58% and FDR of 42%.

In a similar fashion to identifying FRET localizations, we can inversely seek inverse extrema (i.e., minima instead of maxima or vice versa) of different parameter channels to search for localizations of non-FRET contributions. To test the possibilities of identifying FRET and non-FRET localizations, we simulated the endoplasmic reticulum (ER) with ribosomes either in close proximity to ER membrane proteins, thus assumed to be in translation mode and within FRET proximities, or farther away from the ER and out of the FRET dynamic range. For doing so while preserving a realistic spreading of the molecules, we simulated the ground-truth locations of the ER and the ribosomes by following a TEM micrograph taken from the George E. Palade EM Collection (41). We chose the ER membrane to be tagged by the donor molecules, (i.e., by staining the EMC complex (42)), and the ribosomes to be tagged by acceptor molecules.

Fig. 6 shows the ground-truth data and simulation results with both localizations using beforehand local extrema search (i.e., local maxima for parameter channels 1 and 3 and local minima for parameter channels 2 and 4), while Fig. S4 shows the inverse local extrema search (i.e., local minima for parameter channels 1 and 3 and local maxima for parameter channels 2 and 4). The localizations for non-FRET pairs are not as good as for the interacting FRET-pairs, as could be expected, as they signify the lack of signal, with accuracies of 67±32 nm (SD) for non-FRET compared to 21±12 nm (SD) for FRET and with a TPR of ∼18% compared to 43%, respectively. When taking into account only parameter channel 1, i.e. the delay in the acceptor mean fluorescence lifetime, for the reverse search, (Fig. 6, G, H), the results resemble the local maxima search, with accuracy of 34±25 nm and with a TPR of ∼28% (Fig. 6, D, G). Nevertheless, the overall structure of ER with interacting ribosomes overlapping with its membrane with the addition of the non-interacting ribosomes is preserved (Fig. 6, E, H).

**Fig. 6.**
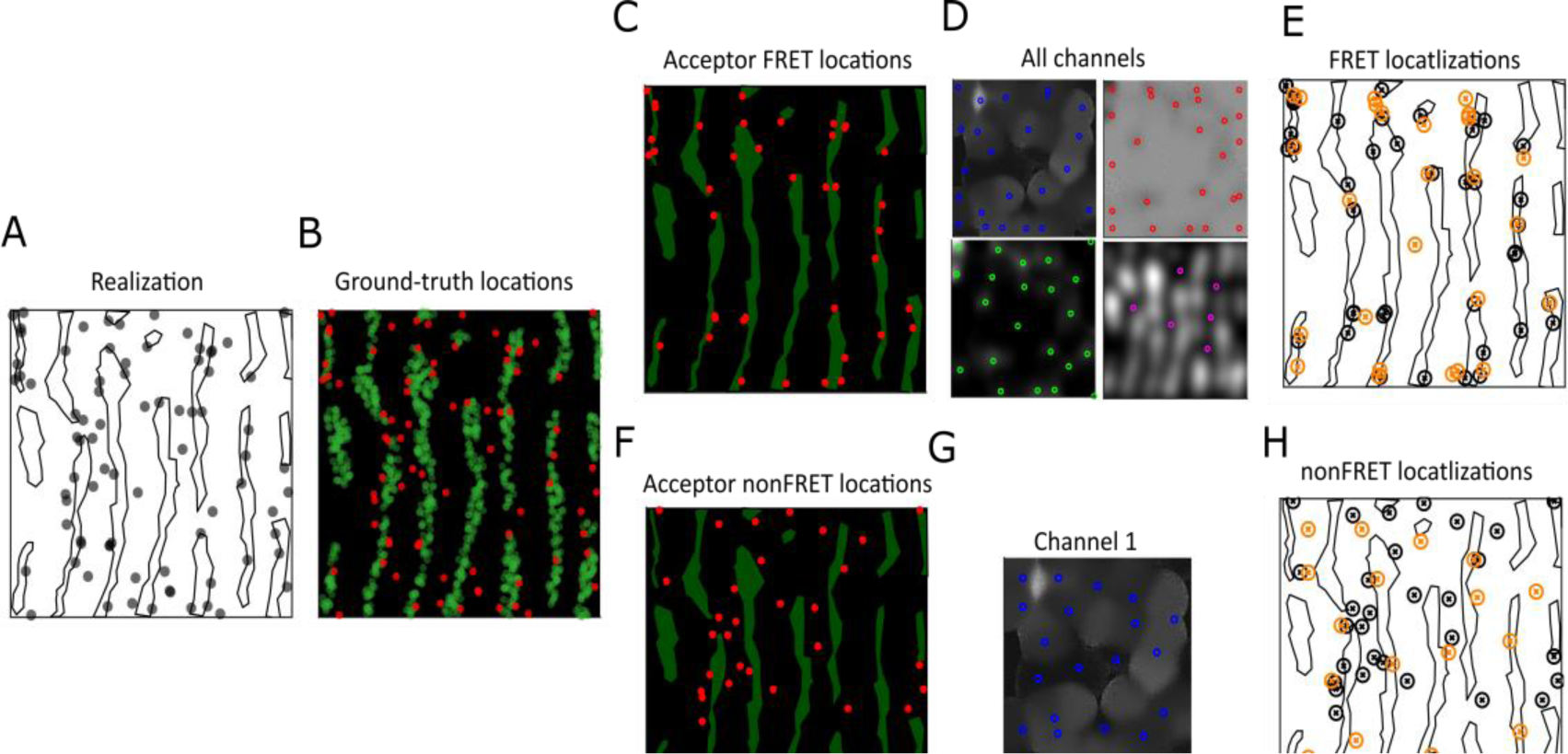
ER-Ribosomes simulation. **A**. Illustration of the micrograph taken to simulate the ER with ribosome molecules. **B**. Donors and acceptors representing ER (#1,100; green dots) and Ribosomes (#120; red dots), respectively, were distributed over a 1×1 µm^2^ frame according to an online available TEM micrograph. **C, F**. The majority of the ribosomes were spread far from the ER (#80; panel **F**), while 33% were spread in close proximity (#40; **C** red dots). **D**. The output of the four parameter channels with localizations of each channel (delay in the acceptor mean fluorescence lifetime in blue; donor mean fluorescence lifetime in red; acceptor to donor fluorescence intensity ratio in green; donor fluorescence intensity in pink). **E**. The ground-truth of interactions represented as black ⊗ signs. Localizations of suspected interacting pairs (orange ⊗ signs) presented next to the ground-truth locations. Comparing localizations with true locations resulted in accuracy of 21±12 nm (SD) with TPR of 43% and FDR of 30% (CSI=0.37). **G**. Parameter channel 1 (delay in the acceptor mean fluorescence lifetime) with localizations of local minima search. **H**. The ground-truth of non-interacting ribosomes represented as black ⊗ signs. Localizations of suspected areas with no interaction (orange ⊗ signed) presented next to the ground-truth locations. Comparing these localizations with true locations yielded detection accuracy of 34±25 nm (SD) with TPR of 28% and the FDR of 42%.

### Testing FRETsael on a well-defined protein-protein interaction *in cellulo*

After testing the strengths and characterizing the advantages and limitations of the FRETsael approach on simulated data, where the ground-truth of each donor– and acceptor-labeled biomolecule is known, we move to testing the capabilities of FRETsael on localizing protein-protein interactions in cells from two-color laser scanning confocal-based fluorescence lifetime imaging (FLIM), with the possibility of FRET. We perform this test using human neuroblastoma cells (SH-SY5Y cells), stained with iFluor 488 conjugated to phalloidin to detect the distribution of F-actin and with Alexa Fluor 594 (AF594)-conjugated fluorescent antibodies binding non-muscle myosin IIA (Fig. 7, A-C). iFluor 488-tagged actin, is the donor and the AF594-tagged non-muscle myosin IIA is the acceptor, and they have a well-known established molecular interaction (43–45), thus rendering them perfect for a FRET-based interaction localization test, especially in the absence of molecular level spatial ground-truth. Importantly, here FP and FN events are not known. The only thing that is known is that FRET events are expected in close proximity to the middle of the actin filaments, obviously controlled by the stoichiometry between the donor and acceptor-labeled antibodies.

**Fig. 7.**
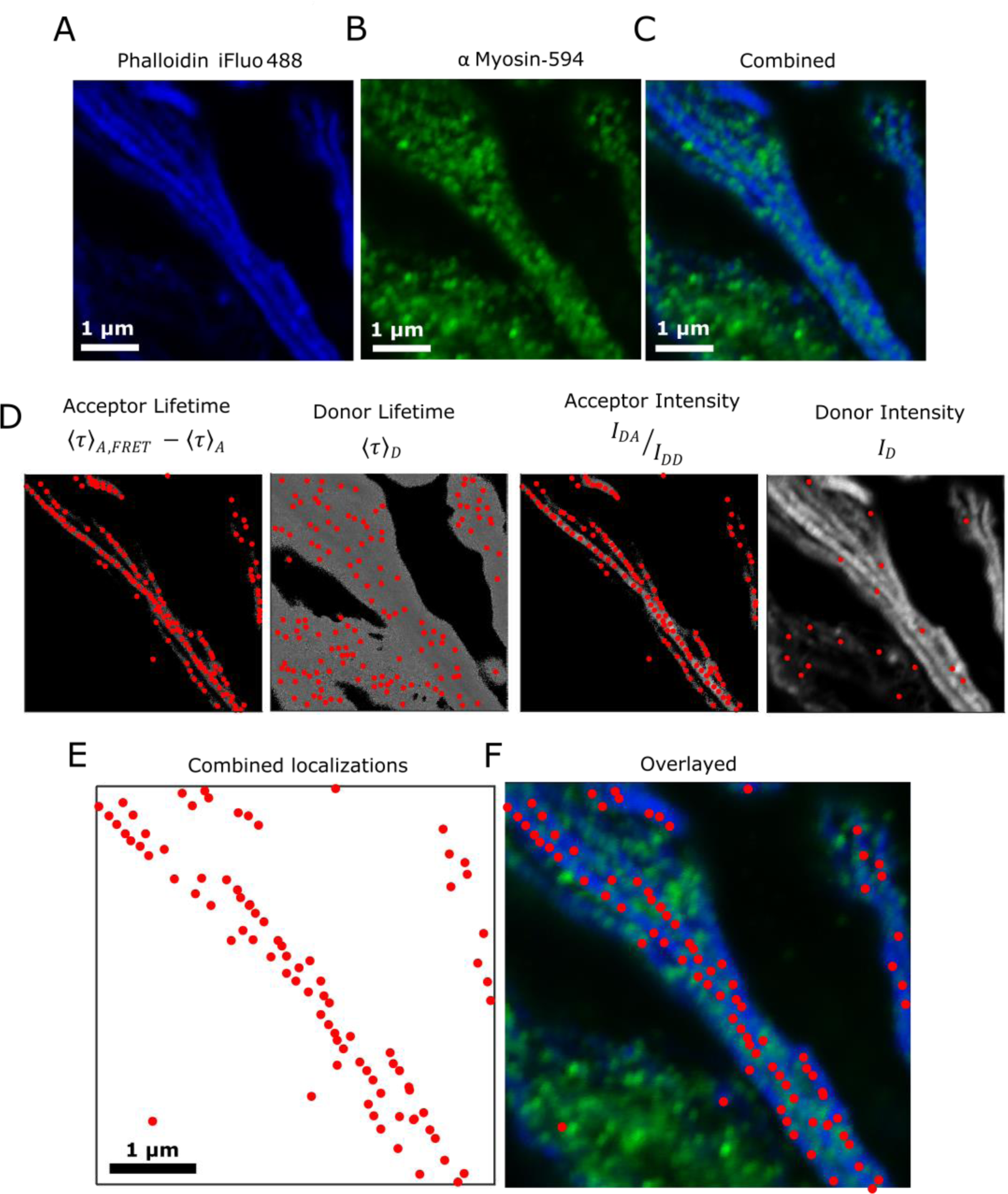
FRET imaging of iFluor 488-phalloidin and AF594-anti-Myosin antibodies in SH-SY5Y cells and FRETsael analyses. **A-B**. Fluorescence intensity images of iFluor 488 conjugated to phalloidin and Alexa Fluor 594 conjugated fluorescent antibodies binding non-muscle myosin IIA. **C**. Merge of panels **A** and **B**. 10×10 µm^2^, 512×512 pixels. Scale bar 1 µm. **D**. Shown from left to right are images of the four different parameter channels with localizations of each channel (delay in the acceptor mean fluorescence lifetime; donor mean fluorescence lifetime; acceptor to donor fluorescence intensity ratio; donor fluorescence intensity). Localizations are depicted in red circles. **E**. localizations after integrating the parameter channels. **F**. Overlay of the merged fluorescence intensity image and localizations extracted after integrating the two optimal channels together.

We performed PIE-based FRETsael experiments of a 10×10 µm^2^ frame with 512×512 pixels, which is achieved using a laser scan step size of ∼20 nm. Importantly, with such small laser scan steps attaining laser scanning confocal microscopy images will end with imaged elements (e.g., actin filaments) that are wider and blurred, owing to the overlap between consecutive pixels. However, this is key for exposing the fine nm-scale differences that we seek. With pixel dwell time of 0.5 ms, the acquisition time of a single frame was ∼130 seconds. Then, we employed the FRETsael analysis on the photon data of the acquired images and report the results (Fig. 7).

Overall, after integrating the four parameter channels, there is a total of 86 detected localizations. Indeed, it seems that the two parameter channels contributing best to the localization are channels 1 (delay in the acceptor mean fluorescence lifetime) and 3 (acceptor to donor fluorescence intensity ratio), while for channels 2 and 4 (donor mean fluorescence lifetime and donor fluorescence intensity respectively) localizations are sparse and spread around the cell. Notably, unlike in the simulations where the donor fluorescence lifetime in the absence of FRET was kept constant, in the experiment it is not promised that the donor will have a constant intrinsic fluorescence lifetime. Therefore, we can rely mostly on the parameters based on the delay in the acceptor mean fluorescence lifetime and the acceptor to donor fluorescence intensity ratio after donor excitation. Accordingly, we utilize harsher conditions for the localizations (higher threshold than in the simulations), when based on parameter channels 2 and 4 (donor mean fluorescence lifetime and donor fluorescence intensity respectively). Based on the localizations found in images of parameter channels 1 and 3, the localizations of potentially-interacting myosin and actin molecules are distributed near the middle of actin filaments. This observation coincides with what Wang *et al.* showed for the organization of the non-muscle myosin in the middle of the actin filament (45).

Importantly, we conducted the same procedure over two controls: (i) one that was stained only for actin (iFluor 488 conjugated to phalloidin), representing the case where there are only donor molecules, and (ii) the other stained with Alexa Fluor 594 (AF594)-conjugated fluorescent antibodies binding non-muscle myosin IIA, to represent an acceptor only scenario (Fig. S5). In both cases, there should be no visible interaction. In the case of donor-only (Fig. S5 A-D), there is a minor leakage from the blue channel into the green channel, however it has no effect for the FRETsael algorithm, where there are localizations only for parameter channels 2 and 4 (donor mean fluorescence lifetime and donor fluorescence intensity respectively). Furthermore, combination of all four parameter channels, tossed aside these localizations as well. In the case of acceptor only (Fig. S5 E-H), none of the parameter channels yielded localizations to begin with.

We tested again the possibility that the effects caused by sparse FRET pairs are blurred due to dense donor only and acceptor only molecules around. We examine this by looking at parameter channel 1, the delay in acceptor mean fluorescence lifetime due to contribution from FRET, in a position where we assume there is a FRET localization (Fig. S6 A, C, E) versus a random location over the cell area (S6 B, D, F). It is clearly observable that the delay is distinct at the FRET position, and monotonically reduced when getting farther away from this position, while in the random position case, the delay is purely stochastic and there is no observable monotonic trend.

We conclude that using FRETsael to analyze PIE-based FRET images using few nm laser scan step sizes can expose FRET localizations with 20-30 nm accuracies and with high sensitivities, and by that reveal the localizations of biomolecular interactions in the cell.

## Discussion

In this work, we have presented the concept of FRET-sensitized acceptor emission localization (FRETsael), in which FRET pairs of interacting biomolecules are localized with 20-30 nm accuracy by finding local extrema of the contribution to FRET in confocal-based PIE FLIM-FRET imaging with few nanometer laser scan step sizes. Importantly, the best experimental parameter channels to retrieve the local maxima of the contributions to FRET rely on FRET-sensitized acceptor emission, rather than the heavily-used reduction in donor fluorescence features (e.g., intensity, lifetime). The work results in the ability to super-resolve biomolecular interactions leading to FRET, without using special tags that blink on-and-off at predefined rates. By using a relatively simple confocal microscopy setup, where the only extensions come from the addition of FLIM with PIE capabilities and using laser scan steps of few nm rather than steps of ∼λ/2.

It is noteworthy that this work focused on FRETsael assuming the donor and acceptor emitters are spread in a given plane, and hence can be retrieved in a 2D scan. This is not necessarily the case in all types of biomolecules imaged in the cell. In practice, the donor– and acceptor-tagged biomolecules could exist at different depths across the Z-axis, while sharing close localizations in the X-Y plane. While the X-Y part of the Gaussian profile of a PSF was used in this work, its profile along the Z-axis is much wider. If the width of the Gaussian PSF profile in the X-Y plane could be ∼200 nm, and in proper confocal setup improved close to 100 nm, across the Z-axis it could be in the range of 600-1,500 nm range, depending on the numerical aperture and the magnification of the objective lens under use.

In this work, we found a limit of X-Y localization accuracy of 20-30 nm, which is ∼1 order of magnitude smaller than the Gaussian PSF profile in the X-Y plane. This region of the Gaussian profile in the X-Y plane is the region where the contribution to fluorescence will be almost similar even if the profile is moved by a few nanometers. Therefore, and in analogy to the profile in the X-Y plane, in the Z-axis, the Gaussian profile with widths in the range 600-1,500 nm could in principle yield a localization accuracy of 60-150 nm in Z. This extension to FRETsael will facilitate interaction localization in 3D. Therefore, in 2D FRETsael, if all donors and acceptors are found at different depths within the 60-150 nm range, their localization using the 2D-scanned images will be sufficient. The only possible difference would be that emitters that are in FRET proximity in the X-Y plain might be far apart in depth, and hence not yield FRET signatures. Nevertheless, if such a pair of dyes appears, it will not exhibit a delay in the acceptor fluorescence decay, and hence will not exhibit much difference between the delay in the acceptor mean fluorescence lifetime after donor excitation versus after acceptor excitation. Yet, in the 3D space within the cell, it is possible that layers of biomolecular interactors will be found at many depths beyond the 60-150 nm range. To employ FRETsael on such biomolecular interactions would require acquiring data in Z-stacks, and only after testing the accuracy and sensitivity of the FRETsael approach in 3D parameter scanning and localizing local extrema.

Here, we show that the most trustworthy parameter among many, for the FRETsael approach, is the difference between the acceptor mean fluorescence lifetime after donor excitation versus after acceptor excitation. It is important, however, to remember that there are situations in which the contribution to FRET is high, but no local extrema can be found using this parameter. If the donor-acceptor distance is short enough relative to the Förster distance, then the delay of the excitation energy at the donor excited-state becomes so short to the level in which it becomes almost undetectable. Inspecting this possibility against the common pairs of fluorescent tags, their majority exhibit Förster distances in the range of 5.0-6.5 nm (46). This, in turn, means that only donor-acceptor distances <3 nm will lead to complete lack of detectability of the biomolecular interactions through this parameter FRET contributions. However, when using FPs, which add their own sizes to the overall biomolecular complex, the distance between the donor and acceptor fluorophores within the fluorescent proteins will be mostly >3 nm. Another common tagging technique is by using primary or secondary fluorescently-labeled antibodies. Since the antibodies also add their own sizes, the potential of having undetectable FRET also becomes improbable. Of course, it is always possible to lower the threshold for detection that is to reduce the minimum reliable value for the delay in order to detect also shorter-length interactions. However, we would rather have a more stringent analysis with sufficiently high thresholds. Lowering thresholds will potentially increase false positive detections. It is a matter of balances as having false negatives, i.e. not detecting a true pair of molecules interacting is less severe than false positive detections, i.e. assuming there is an interaction while there is none.

Overall, the ability to localize biomolecules with spatial accuracies similar to the biomolecular sizes helps understand where biomolecules are accumulating in the cell, whether they cluster together or not, and all in the context of cellular compartments, ranging from optically resolvable µm size, down to difficult or even impossible to optically resolve nm size. Therefore, FRETsael could elucidate unknown features of protein-protein interactions such as those existing in nm-sized compartments and other entities, such as the mitochondrial intermembrane space (47, 48), within endoplasmic reticulum tubes (49, 50), in nuclear nanodomains (51, 52), in autophagy (53, 54) or in stress granules (55, 56). These examples and others are potential targets for implementing FRETsael to study how specific biomolecules interact, where within these cellular and sub-cellular domains, and perhaps even how they change in response to different stimuli.

These concepts are in general at the heart of SR light microscopy. SR techniques do not overcome Abbe’s diffraction limit, but rather bypass it via acquiring molecular contrast that can be used to get refined localizations. While in SMLM the molecular contrast is acquired via bright and dark states of special fluorophores, in STED the molecular contrast is acquired via regions where emission occurs immediately as the depletion laser reaches the sample through stimulated emission and other regions with spontaneous emission. We show the achievement of this molecular contrast in figures S3 and S6; while a FRET pair will result in a long delay in the acceptor mean fluorescence lifetime due to contribution from FRET, which will decrease when getting farther away from the center position of the pair, a donor only position will exhibit no such effect, and the delay, if any, is constant throughout the change of the distance from the center position of the donor.

FRETsael serves as the basis of yet another SR localization imaging technique that builds on using the non-uniformity of the excitation profile. In many ways, FRETsael carries with it analogies to MINFLUX, where acquiring data from moving the approximately paraboloid profile formed within the doughnut-shaped PSF relative to the sample is used for localizing the position of an emitter. In FRETsael, we use the knowledge of the shape of the PSF, this time a Gaussian, to localize a FRET-pair rather than a single emitter. The main difference between MINFLUX and FRETsael is that while in MINFLUX the molecular contrast is gained by changes between brighter regions than others of a given emitter relative to its position within the paraboloid of the doughnut-shaped PSF, in FRETsael the molecular contrast is achieved via changes between regions with FRET contributions and others with non-FRET contributions (Fig S5, S6).

Regarding the matter of acquisition speed, one major drawback of FRETsael in its current version is the slow acquisition time. To attain trustworthy pixel-based mean fluorescence lifetime data, 0.5 ms pixel dwell time can be considered, and then if a 10×10 µm^2^ frame is scanned using 512×512 pixels (i.e., ∼20 nm laser scan size), one frame is attained after ∼130 seconds. Such slow acquisition times render FRETsael useful mostly for fixed cells and slow processes in live cells. Acquisition times in FRET imaging could be improved if wide-field microscopes would be used instead of a laser scanning confocal-based microscope. Indeed, fast cameras, EMCCD or sCMOS, exhibit integration times of a few ms. However, most fast cameras do not exhibit pixel-based fluorescence lifetime capabilities. Single-photon avalanche diode (SPAD)-based arrays are now emerging, where each pixel can have lifetime capabilities (57–60). We envision the next generation of FRETsael would be in the context of a wide-field microscope with lifetime-based SPAD arrays acting as lifetime-capable cameras, which will allow nm localizations of biomolecular interactions in live cells in as close as it can get to the biological equivalent of real-time.

## Online Methods

### Known ground-truth confocal-based imaging simulations

We positioned donors and acceptors in an X-Y plane, with varying (a) donor-acceptor distances, (b) densities, (c) donor:acceptor stoichiometries, (d) laser scan step sizes, and (e) widths of the Gaussian excitation profile. Then, we performed Gaussian-shaped PSF blurring for both the donors and acceptors if they were simulated as contributing fluorescence either from FRET-sensitization or from non-FRET contributions. Then, we simulated the respective photon detection times, both macrotimes (i.e., the absolute photon detection times) and nanotimes (i.e., the photon detection times relative to the moment of excitation that led to their emission), using MC calculations, for three different photon streams: (i) donor fluorescence photons following donor excitation, (ii) acceptor fluorescence photons following donor excitation and FRET, and (iii) acceptor fluorescence photons following acceptor excitation. These photon streams are the streams in use in PIE-FRET experiments. Therefore, in case of donor excitation, the possibility of direct acceptor excitation with examined against a direct acceptor excitation factor, *dir*, where if true, a non-FRET acceptor photon following donor excitation was sampled, however, if false, donor excitation was examined. In the next step, if FRET from this donor to a nearby acceptor was feasible, the choice of whether it was a donor or an acceptor photon that was emitted, was evaluated against the FRET efficiency value for this donor-acceptor pair (eq. S21). In the case that FRET occurred, the emitted photon was recorded as an acceptor photon following donor excitation. If FRET did not occur, donor fluorescence leakage to the acceptor channel was also checked, against a leakage factor, *lk*. Only if FRET and leakage did not occur, was the photon recorded as a donor fluorescence after donor excitation photon. As for the photon nanotimes, we sampled their values from exponential distributions, with means as the donor and acceptor fluorescence lifetimes in the absence of FRET (eqs. S17, S18), or due to FRET (eqs. S13, S14), considering the relevant donor-acceptor distances, driven from FRET theory equations (see Supplementary Text). Importantly, these simulations assume that either a donor or an acceptor that are not involved in FRET will have unchanging fluorescence lifetimes, an assumption that is ideal relative to actual FLIM results. For simplicity we used the literature parameters of eGFP and mCherry as a common FP-based FRET pair (36, 37). We weighted the fluorescence contributions by the position of the biomolecular emitters relative to the Gaussian profile of the PSF at each given scanning position, the result of which is mathematically described in the supplementary text (eqs. S19, S20). Then, we iterated these simulations, after shifting the Gaussian-shaped PSF in X and Y, in a fashion similar to nm laser scanning steps we later performed in actual acquisitions of laser scanning confocal-based images (Fig. S1). The results of the simulations are fluorescence intensity and mean fluorescence lifetime images in the three-photon streams. Then, these images are used for calculating images of the parameter channels (see below), which allow the analyses of localizations of highest contributions to FRET.

### FRETsael analysis

The FRETsael algorithm uses built-in *Matlab* functions in the following manner. The FLIM input data includes both the photon detection event macrotimes and nanotimes. The number of photon detection macrotimes and the mean of photon nanotimes per pixel in the scanning area were used for calculating the pixel intensities and mean fluorescence lifetimes, respectively, per each detection channel, where the mean photon nanotime is equivalent to the intrinsic mean fluorescence lifetime (eqs. S29-S31). Then, we calculate images of the parameter channels (see eqs. 1-4, and their relation to FRET in eqs. S21, S22-S24, S25, and S26). In the next step, we apply cross-correlation between the relevant image for each parameter channel and a Gaussian intensity profile for the laser PSF. Next, we apply a local maxima search algorithm to find possible locations for FRET pair position. Note, for parameter channels 2 and 4, where FRET is demonstrated with reduction in the lifetime or intensity, we apply the cross-correlation with the inverse image. At that point, we have possible localizations for each of the four-parameter channels. The final step is to filter out localizations that only appear in one of the parameter channels. Thus, a localization is considered true only if there is another localization in a different parameter channel within 50 nm proximity.

### Experimental setup

We performed fluorescence lifetime imaging (FLIM)-FRET (FLIM-FRET) using a confocal-based microscopy setup (ISS^TM^, Champaign, IL, USA) assembled on top of an Olympus IX73 inverted microscope stand (Olympus, Tokyo, Japan). We used PIE with 532±1 and 488±1 nm pulsed picosecond lasers (FL-532-PICO, CNI, China and QuixX® 488-60 PS, Omicron, GmbH; pulse width of 100 ps FWHM, operating at 20 MHz repetition rate and 100 or 40 μW, respectively, measured at the back aperture of the objective lens) for exciting Alexa Fluor 594-antibodies binding non-muscle myosin IIA and iFluo488-phalloidin-F-actin, respectively. The laser beam passes through a polarization-maintaining optical fiber (P1-405BPM-FC-Custom, with specifications similar to those of PM-S405-XP, Thorlabs, Newton, NJ, USA) and after passing through a collimating lens (AC080-016-A-ML, Thorlabs), the beam is further shaped by a quarter waveplate (WPMP2-20(OD)-BB 550 nm, Karl Lambrecht Corp., Chicago, IL, USA) and a linear polarizer (DPM-100-VIS, Meadowlark Optics, Frederick, CO, USA). A major dichroic mirror (DM) with high reflectivity at 488 and 532 nm (ZT405/488/532/640rpc-XT, Chroma, Bellows Falls, Vermont, USA), for iFluor 488 and Alexa Fluor 594, reflects the light to the optical path through galvo-scanning mirrors (6215H XY, Novanta Corp., Boston, MA, USA) and scan lens (30 mm Dia. x 50 mm FL, VIS-NIR Coated, Achromatic Lens, Edmund Optics, Barrington, NJ, USA; both used to acquire the scanned image), and then into the side port of the microscope body through its tube lens, positioning it at the back aperture of a high numerical aperture (NA) super apochromatic objective (UPLSAPO100XO, 100X, NA=1.4, oil immersion, Olympus), which focuses the light onto a small effective excitation volume, positioned within the sample chamber (µ-Slide 8 Well high Glass Bottom, Ibidi, Gräfelfing, GmbH). Scattered light returns in the excitation path, and a fraction of it is imaged on a CCD camera, used Z-positioning, using Airy ring pattern visualization. Fluorescence from the sample is collected through the same objective lens, is transmitted through the major DM and is focused with an achromatic lens (25 mm Dia. x 100 mm FL, VIS-NIR Coated, Edmund Optics) onto a 100 μm diameter pinhole (variable pinhole, motorized, tunable from 20 μm to 1 mm, custom made by ISS^TM^), and then re-collimated with another achromatic lens (AC254-060-A, Thorlabs). Fluorescence is then split between two detection channels, the acceptor and donor detection channels, using a DM with a cutoff wavelength at λ=605 nm for iFluor 488 and Alexa Fluor 594 (FF605-Di02-25×36, Semrock Rochester, NY, USA). Then, the fluorescence is further cleaned using a 615/24 nm, for Alexa Fluor 594, and 510/20 nm, for iFluor 488, single band bandpass filter (FF01-615/24-25, or FF01-510/20-25, Semrock, Rochester, NY, USA). Fluorescence was collected using two hybrid PMTs (R10467U-40, Hamamatsu Photonics, Shizuoka, Japan), routed to a time-correlated single photon counting (TCSPC) card (SPC 150N, Becker & Hickl, GmbH). Images were attained by using a laser scanning module (LSM), in which a 3-axis DAC module (custom made by ISS^TM^) synchronized the data acquisition and control over the X and Y galvo-scanning mirrors, which assisted in bringing the effective excitation volume to different positions to acquire pixel data per a given Z layer (see Fig. 1, A for schematics of the laser scanning confocal-based FLIM setup with PIE capabilities)

The scanning conditions are: (i) pixel dwell time – 0.5 ms; (ii) number of pixels 512×512; (iii) field of view area –10×10 µm^2^; (iv) number of acquired frames per image is 5. Data acquisition is performed using the VistaVision software (version 4.2.095, 64-bit, ISS^TM^) in the time-tagged time-resolved (TTTR) file format. In FLIM, images were attained by calculating the mean photon nanotime for all photons acquired in a given pixel per detection channel and photon stream. Calculation mean photon nanotimes were performed on pixels only if they had >150 photons of the given photon stream.

### Cell work

#### Cell cultures

Human-derived SH-SY5Y neuroblastoma cells were cultured in a complete medium (DMEM/F12, 10% FBS, 1% L-glutamine, 1% Pen-Strep, 1% sodium pyruvate, 1% NEAA) and incubated at 37°C and in 5% CO2. Cells were sub-cultured once a week by trypsin, half of the medium was changed four days after passaging. Two days before conducting the experiment, the cells were transported to grow in an 8 well µ-Slide (Ibidi, Gräfelfing, GmbH).

#### Protocol for immunofluorescence with formaldehyde fixation

The cells’ media was washed with PBS, followed by a 15-minute incubation in 4% formaldehyde in PBS solution at room temperature, and then performing three more washes with PBS. To enhance cellular permeability, a 10-minute treatment with 0.1% Triton X-100 in PBS is conducted, followed by three washes with PBS. Blocking unspecific binding sites was achieved by incubating the cells in a 1% BSA PBS solution for 30 minutes, followed by three PBS washes. Next, the cells are exposed to 0.1 mg/mL of a primary antibody against myosin (anti-non-muscle Myosin IIA antibodies, ab138498, Abcam) in 1% BSA for an hour. Post-incubation, three washes with PBS are performed to remove excess unbound antibodies. Subsequently, a one-hour incubation in the dark with the secondary antibody (goat anti-rabbit IgG H&L, labeled with Alexa Fluor 594, ab150080, Abcam) in 1% BSA, alongside staining actin with phalloidin-iFluor 488 (ab176753, Abcam) at a 1:1000 dilution, is conducted. Finally, the cells undergo three PBS washes to ensure the removal of excess reagents.

## Data availability

The raw PIE-FLIM data of actin-myosin measurements is provided over Zenodo: https://doi.org/10.5281/zenodo.10283490.

## Code availability

The *Matlab* scripts for the simulations and analyses are provided over Zenodo: https://doi.org/10.5281/zenodo.10283490.

## Author Contributions

E.L. conceived the idea and concept behind FRETsael and developed the theory behind it, Y.R. developed the algorithms and code for simulating confocal-based images and for analyzing FRET localizations using FRETsael. Y.R. performed all simulations. Y.R., P.D. and S.K. performed all cell preparations. Y.R. and P.D. analyzed the FLIM-FRET imaging data using the FRETsael code. E.L. secured funding for this project. Y.R. and E.L. wrote the initial version of this article and all authors contributed to finalizing the writing of the article.

## Acknowledgements

This work was supported by the Israel Science Foundation (ISF, grant 556/22 to E.L.) and the European Union’s Horizon Europe Research and Innovation Programme under the EIC Pathfinder-Open grant agreement #101099654 (RT-SuperES to E.M.). E.M. is the Arthur Gutterman Family chair for Stem Cell Research.

