## Supporting Materials for "FRET-sensitized acceptor emission localization (FRETsael) – nanometer localization of biomolecular interactions using fluorescence lifetime imaging"

Supplementary Material

### <u>Supplementary Text</u>

<u>Theory supporting the parameter channels used in FRETsael</u>

Following the explanation of the FRETsael concept, we put it in the proper theoretical context. Assume a system of two dyes, donor ($D$ ground state and $D^*$ excited state) and acceptor ($A$ ground state and $A^*$ excited state), in which (i) the donor can be directly excited, and (ii) the donor de-excitation energy levels overlap with the acceptor excitation energy levels; hence the donor fluorescence spectrum overlaps with the acceptor excitation spectrum. If the distance between $D$ and $A$ is too large (typically >10 nm), the time evolution of the donor excited-state can be described by an exponential decay with a de-excitation rate, $k_{F,D}$, after it was excited at $t$=0 (eqs. S1, S2).

$$D^* \overset{k_{F,D}}{\rightarrow} D + h\nu_D + \Delta \quad \text{(Eq. S1)}$$

$$\frac{dD^*}{dt} = -k_{F,D}D^* \quad \text{(Eq. S2)}$$

where $k_{F,D}$ is the rate of de-excitation of the donor $D^* \rightarrow D$, $h\nu_D$ is radiative energy with donor fluorescence wavelengths and $\Delta$ is non-radiative energy. Eq. S2 describes the evolution of the donor excited-state as a function of time following excitation, assuming this is the only de-excitation process. The solution to it is a mono-exponential decay, with a rate, $k_{F,D}$.

In the ideal case, in which the acceptor dye is not excitable with the donor excitation wavelength, the acceptor stays at ground state. If the distance between $D$ and $A$ is not too long (typically <10 nm), the excitation energy can also be depleted from the donor due to radiation-less transfer of the excitation energy to the acceptor, with a rate constant $k_{FRET}$ (eq. S3). This, in turn, can be added as another donor de-excitation process (eq. S4).

$$D^* + A \overset{k_{FRET}}{\rightarrow} D + A^* \overset{k_{F,A}}{\rightarrow} D + A + h\nu_A + \Delta \quad \text{(Eq. S3)}$$

where $k_{FRET}$ is the rate of donor-acceptor excitation energy transfer, $k_{F,A}$ is the rate of de-excitation of the acceptor $A^* \rightarrow A$, and $h\nu_A$ is the radiative energy with acceptor fluorescence wavelengths. This results in: (i) another donor de-excitation route, and (ii) the excitation of the acceptor. Following this, the time evolution of the donor and acceptor excited states can be described (eqs. S4, S5, respectively).

$$\frac{dD^*}{dt} = -\left(k_{F,D} + k_{FRET}\right)D^* \quad \text{(Eq. S4)}$$

$$\frac{dA^*}{dt} = k_{FRET}D^* - k_{F,A}A^* \quad \text{(Eq. S5)}$$

It is important to mention that $k_{FRET}$ is different for different donor-acceptor distances, $R_{DA}$, and for different donor and acceptor spectroscopic characteristics, expressed in $R_0$, the Förster distance, as is shown in eq. S6.

$$k_{FRET}(R_{DA}) = k_{F,D}\left(\frac{R_0}{R_{DA}}\right)^6 \quad \text{(Eq. S6)}$$

where the Förster distance is dependent on multiple factors (eq. S7).

$$R_0 = 0.2108\left(\frac{\kappa^2 \Phi_{F,D(0)}}{n_{im}^4}\int \overline{F_D}(\lambda)\varepsilon_A(\lambda)\lambda^4 d\lambda\right)^{1/6} \quad \text{(Eq. S7)}$$

such as $\Phi_{F,D(0)}$, the donor fluorescence quantum yield in the absence of an acceptor, the integrand which represents the donor fluorescence spectrum in the absence of an acceptor, $\overline{F_D}(\lambda)$, and the acceptor excitation spectrum with extinction coefficient, $\varepsilon_A(\lambda)$ (in $M^{-1}cm^{-1}$), at the wavelength λ (in nm), $\kappa^2$ as the orientation factor quantifying the relative orientation of the dipoles of the donor and acceptor dyes, and $n_{im}$ which is the local refractive index between the donor and acceptor dyes.

Additionally, the quantum yield of the FRET-based donor deexcitation process, better known as the FRET efficiency, E, is given by (eq. S8)

$$E = \frac{k_{FRET}}{k_{F,D}+k_{FRET}} = \frac{1}{1+\left(\frac{R_{DA}}{R_0}\right)^6} \quad \text{(Eq. S8)}$$

The outcome of eqs. S6-S8 is that for a single given $R_{DA}$, a given FRET rate and a given FRET efficiency exists. Obviously in reality, a distribution of distances exists, and hence each distance within the distribution will contribute a fraction of the value of the distribution with its own de-excitation rate to the overall FRET and fluorescence. For the sake of simplicity, we will stay within the approximation of an average $R_{DA}$ value. Combining eq. S6-S8 into eqs. S4 and S5, we get (eqs. S9 and S10),

$$\frac{dD^*}{dt} = -\frac{k_{F,D}}{1-E}D^* \quad \text{(Eq. S9)}$$

$$\frac{dA^*}{dt} = \frac{k_{F,D} \cdot E}{1-E}D^* - k_{F,A}A^* \quad \text{(Eq. S10)}$$

The solutions of eqs. S9 and S10 are (eqs. S11 and S12),

$$D^*(t) = D^*_{t=0}e^{-\frac{k_{F,D}}{1-E}t} \quad \text{(Eq. S11)}$$

$$A^*(t) = \frac{D^*_{t=0}k_{FRET}}{k_{FRET}+k_{F,D}-k_{F,A}}\left[e^{-k_{F,A}t} - e^{-(k_{FRET}+k_{F,D})t}\right] = \frac{D^*_{t=0}}{\frac{1}{E}\left(1-\frac{k_{F,A}}{k_{F,D}}\right)+\frac{k_{F,A}}{k_{F,D}}}\left\{e^{-k_{F,A}t} - e^{-\frac{k_{F,D}}{1-E}t}\right\} \quad \text{(Eq. S12)}$$

Instead of $k_{F,D}$ and $k_{F,A}$, it is possible to use fluorescence lifetimes, $\frac{1}{\tau_D}$ and $\frac{1}{\tau_A}$, respectively (eqs. S13 and S14).

$$D^*(t) = D^*_{t=0}e^{-\frac{t}{\tau_D(1-E)}} \quad \text{(Eq. S13)}$$

$$A^*(t) = \frac{D^*_{t=0}}{\frac{1}{E}\left[1-\frac{\tau_D}{\tau_A}(1-E)\right]}\left[e^{-\frac{t}{\tau_A}} - e^{-\frac{t}{\tau_D(1-E)}}\right] \quad \text{(Eq. S14)}$$

While eq. S13 represents a decay, eq. S14 represents a function with an exponential rise, due to the population of acceptor-excited state as a function of FRET rates, and a decay, for the acceptor de-excitation. Therefore, the acceptor de-excitation is delayed, relative to what it could have been if it was directly excited. Additionally, note that the longer the distance is, the longer the delay will be, but the lower the decay amplitude will be (i.e., the value of the pre-exponential factor).

Eq. S14 is an idealized solution, as it does not consider the realistic possibility that a fraction, *dir*, of the donor excitation energy will be invested in the direct excitation of the acceptor. This depends on the ratio of extinction coefficients of the donor and the acceptor at the excitation wavelength intended primarily for donor excitation. Eq. S15 describes the more realistic situation.

$$A^*(t) = \left[\frac{(1-dir)D^*_{t=0}}{\frac{1}{E}\left[1-\frac{\tau_D}{\tau_A}(1-E)\right]} + dir \cdot D^*_{t=0}\right] e^{-\frac{t}{\tau_A}} - \left[\frac{(1-dir)D^*_{t=0}}{\frac{1}{E}\left[1-\frac{\tau_D}{\tau_A}(1-E)\right]}\right] e^{-\frac{t}{\tau_D(1-E)}} \quad \text{(Eq. S15)}$$

When comparing eqs. S14 and S15, it becomes clear that the more acceptor direct excitation occurs at the wavelength intended primarily for donor excitation (i.e., the higher *dir* is), the faster the rise time of the acceptor fluorescence decay will be reached, which should shorten the decay rise time as well as the overall mean lifetime of the decay. Yet, for a given acceptor (acceptor-only or within donor-acceptor pair), and a given donor excitation wavelength, the *dir* factor is fixed.

Additionally, another realization that should be considered is that the majority of fluorescent dyes have extremely long tails of fluorescence spectra, some of which enter the spectral range that is dedicated for the acceptor fluorescence detection. One can say that for a given optical setup and pair of dyes, a donor fluorescence leakage factor, *lk*, which is proportional to the fraction of the donor fluorescence spectrum area that enters the spectral range for acceptor fluorescence detection, is the factor by which donor fluorescence is added to the data collected from the acceptor detection channel. Therefore, the acceptor fluorescence decay can be written to also take this into account (eq. S16).

$$A^*(t) = (1-lk)\left\{\left[\frac{(1-dir)D^*_{t=0}}{\frac{1}{E}\left[1-\frac{\tau_D}{\tau_A}(1-E)\right]} + dir \cdot D^*_{t=0}\right] e^{-\frac{t}{\tau_A}} - \left[\frac{(1-dir)D^*_{t=0}}{\frac{1}{E}\left[1-\frac{\tau_D}{\tau_A}(1-E)\right]}\right] e^{-\frac{t}{\tau_D(1-E)}}\right\} + lkD^*_{t=0}e^{-\frac{t}{\tau_D(1-E)}} \quad \text{(Eq. S16)}$$

Eq. S16 exhibits another shortening of the rise time of the acceptor decay, now due to the additional contributions from the donor fluorescence decay. Yet, like for *dir*, also the effect of *lk* is fixed for a given donor. Notably, if FRET experiments are properly planned, the factors *lk* and *dir* are expected to be small (<10% combined), which should diminish their effect on shortening the delayed acceptor fluorescence decay. Therefore, for the sake of clarity, we will continue to work with the idealized case presented in eq. S14.

Until this point, we described the acceptor fluorescence decay as if all of the molecules are donor-acceptor pairs. In FRET imaging of cells this is not necessarily a given. While a donor-acceptor pair can be ideally described with eq. S14, other molecules in the field of excitation that have donors with no nearby acceptor, or acceptor with no nearby donors may contribute non-FRET contributions to the acceptor detection channel (eqs. S17 and S18, respectively).

$$D^*(t) = lk \cdot (1 - dir) D^*_{t=0} e^{-\frac{t}{\tau_D}} \quad \text{(Eq. S17)}$$

$$A^*(t) = dir A^*_{t=0} e^{-\frac{t}{\tau_A}} \quad \text{(Eq. S18)}$$

The non-FRET contributions in eqs. S17 and S18 are per each donor or acceptor that are not within FRET distances. Now, assume an illuminated region of a diffraction-limited approximate Gaussian PSF has various donors not in FRET, $n_D$, various acceptors not in FRET, $n_A$, and various donor-acceptor pairs, $n_{FRET}$, found at different locations along the X-Y plane. Additionally, assume the function *psf(x,y)* (in 2D, in this work) describes the Gaussian distribution of excitations, and hence eventually of contributions to photon rates. Then, one can describe the overall contribution to the acceptor fluorescence decay as eq. S19.

$$A^*{}_{overall}(t) = \sum_{i=1}^{n_D} psf(x_i, y_i) \cdot \left[ lk \cdot (1 - dir) D^*_{t=0} e^{-\frac{t}{\tau_D}} \right] + \sum_{i=1}^{n_A} psf(x_i, y_i) \cdot \left( dir A^*_{t=0} e^{-\frac{t}{\tau_A}} \right) + \sum_{i=1}^{n_{FRET}} psf(x_i, y_i) \cdot \left\{ \frac{D^*_{t=0}}{\frac{1}{E}\left[1 - \frac{\tau_D}{\tau_A}(1-E)\right]} \left[ e^{-\frac{t}{\tau_A}} - e^{-\frac{t}{\tau_D(1-E)}} \right] \right\} \quad \text{(Eq. S19)}$$

Therefore, obviously, the higher the fraction of donor-acceptor pairs that undergo FRET is over the number of molecules in the field of view and the more donor-acceptor pairs are closer to the center of the Gaussian-shaped excitation profile (the higher the value of $psf(x_i, y_i)$ is), the larger its contribution will be to the overall signal and hence more delayed the overall acceptor decay, $A^*{}_{overall}(t)$, will be. One also has to consider the basic prerequisite that the molecules at hand are usually proteins fused to fluorescent proteins (FPs), or proteins tagged with fluorescently-labeled antibodies, and hence each such "labeled" protein takes a self-size of a few nm. Therefore, it is plausible to believe that at a given 10x10 $nm^2$ area only a few molecules are expected, and that these donor-acceptor molecules will either be in FRET distances (<10 nm) or not. Then, one can take the Gaussian PSF profile into account and can assume that if a donor-acceptor pair is positioned at the maximum of the PSF, and non-FRET donor and acceptor molecules are positioned at other regions away from it, on the periphery of the Gaussian PSF, the contribution to FRET will be higher than the contribution to non-FRET, and hence the delay of the acceptor decay will be maximized. Nevertheless, this also depends on the ratio of donor-acceptor pairs at the center of the PSF to the non-FRET donors and acceptors on the periphery of the PSF. Therefore, the maximization of the delay of the acceptor fluorescence decay will always be relative and not absolute.

These derivations, of course, are idealized to the case of a single donor to single acceptor one-to-one FRET. One-to-many and many-to-one FRET pathways may occur, but within the 10x10 $nm^2$ assumption are not expected to contribute much. That is true only in 2D. In 3D, many more molecules can co-exist (e.g., in a 10x10x40 $nm^3$ voxel).

As an alternative in this context, the overall donor fluorescence decay can also be used (eq. S20).

$$D^{*}{}_{overall}(t,x,y) = \sum_{i=1}^{n_D} psf(x_i,y_i) \cdot \left(D_{t=0}^{*} e^{-\frac{t}{\tau_D}}\right) + \sum_{i=1}^{n_{FRET}} psf(x_i,y_i) \cdot \left[D_{t=0}^{*} e^{-\frac{t}{\tau_D(1-E)}}\right] \quad \text{(Eq. S20)}$$

where one expects to examine the decrease in donor lifetime. However, other sources of donor lifetime reductions can come from micro-environmental quenching. Therefore, while the decrease in the donor lifetime might not be sensitive enough, the delay of the acceptor fluorescence decay should be more sensitive, especially when tested against the acceptor fluorescence decay after it has been excited directly (i.e., not via the donor and FRET pathway).

In summary, the following parameters are to be tested when testing the sensitivity of FRETsael: (i) fraction of non-FRET donors/acceptors, (ii) positions of molecules in space relative to the PSF, (iii) densities of molecules, (iv) different donor and acceptor intrinsic lifetimes, and (v) different *lk* and *dir* factors, out of which our simulations tested (i)-(iii).

Importantly, when discussing FRET in time-resolved terms, one has to consider the reduction in donor fluorescence lifetime due to FRET (eq. S21).

$$\tau_{DD} = \tau_D(1-E) \quad \text{(Eq. S21)}$$

where $\tau_{DD}$ is the donor fluorescence decay (after donor excitation) in the presence of an acceptor within FRET distances (typically <10 nm). This value dictates, for a donor excitation event, what is the probability that the de-excitation will occur due to transfer of the energy to the acceptor via FRET, which is at a given donor-acceptor distance, $R_{DA}$. Importantly, in a given image pixel, with contributions from both donor-acceptor FRET pairs and from donor-only molecules, assuming the FRET pair maintains a constant FRET efficiency and the donor fluorescence lifetime in the absence of FRET stays unchanged, pixels with minimal donor fluorescence lifetimes are hence expected to report maximal contributions to FRET.

The above expressions of fluorescence decays include the coefficients $D_{t=0}^{*}$ and $A_{t=0}^{*}$, which are pre-exponential factors that depend on the concentrations, excitation efficiencies and fluorescence quantum yields of the donor and acceptor, respectively. Out of these dependencies, the only parameters that predominantly vary are the donor and acceptor concentrations. Therefore, ratios of these parameters are proportional to the stoichiometries.

Using the above derivations, we can now derive the dependencies of the parameter channels used in this work on FRET as well as other factors.

The delay in the acceptor mean fluorescence lifetime due to FRET:

Take a pure FRET contribution, such as that represented by eq. S14. Let us calculate the acceptor mean fluorescence lifetime after FRET (see eq. S22).

$$\langle\tau\rangle_{DA} = \frac{\int_0^\infty tA^*(t)dt}{\int_0^\infty A^*(t)dt} = \cdots = \tau_A + \frac{\tau_D}{\left[1+\left(\frac{R_0}{R_{DA}}\right)^6\right]} = \tau_A + \tau_D(1-E) \quad \text{(Eq. S22)}$$

where the acceptor mean fluorescence lifetime after FRET decreases monotonically as the FRET efficiency increases. In practice, the mean fluorescence lifetime in the acceptor detection channel is a weighted average of contributions to FRET, and contributions from direct acceptor excitation and from donor fluorescence leakage to the acceptor channel (eq. S23).

$$\langle\tau\rangle_{DA} = f_3\tau_{DD} + f_2\tau_A + f_1[\tau_A + \tau_D(1-E)] \quad \text{(Eq. S23)}$$

where an exact description of the weights of these contributions is given by eq. S19. This shows that the process is a trial to maximize the contribution to FRET, and hence minimize the non-FRET contributions. The practice is to take the difference of the mean fluorescence lifetime of the acceptor after donor excitation, $\langle\tau\rangle_{DA}$, and after acceptor excitation, $\langle\tau\rangle_{AA}$, which is $\tau_A$. If we assume that the contribution from donor fluorescence leakage to the acceptor detection channel, $f_3$, is negligible, we show that the retrieved acceptor mean fluorescence lifetime difference between after donor versus after acceptor excitation is directly related to the FRET efficiency, and to the fraction of the contribution to FRET (eq. S24). If the FRET efficiency values in the system can be described by a distribution with a single central population, then a constant ensemble-averaged FRET efficiency can be assumed, which leaves the contribution to FRET, $f_1$, as the parameter to be maximized.

$$\langle\tau\rangle_{DA} - \langle\tau\rangle_{AA} = f_3\tau_{DD} + f_2\tau_A + f_1[\tau_A + \tau_D(1-E)] - \tau_A = f_3\tau_{DD} + (f_1+f_2)\tau_A + f_1\tau_D(1-E) - \tau_A \approx \tau_A + f_1\tau_D(1-E) - \tau_A = f_1\tau_D(1-E)$$

(Eq. S24)

where $\langle\tau\rangle_{DA}$ and $\langle\tau\rangle_{AA}$ are the acceptor mean fluorescence lifetimes after donor excitation and after acceptor excitation, respectively.

Therefore, in a situation where the acceptor fluorescence decay is a superposition of contributions from FRET and contributions from direct acceptor excitations, the mean fluorescence lifetime will always be a weighted average that will scale from $\tau_A$ and to the value in eq. S22.

<u>Fluorescence intensities and their ratios:</u>

Take a pure FRET contribution and let us calculate the expected donor and acceptor fluorescence intensities after donor excitation, for a pure contribution to FRET (eqs. S25, S26)

$$I_{DD} = \int_0^{\infty} D^*_{t=0} e^{-\frac{t}{\tau_D(1-E)}} dt = \cdots = D^*_{t=0}\tau_D(1-E) \quad \text{(Eq. S25)}$$

$$I_{DA} = \frac{D^*_{t=0}}{\frac{1}{E}\left[1-\frac{\tau_D}{\tau_A}(1-E)\right]}\left[\int_0^{\infty} e^{-\frac{t}{\tau_A}} dt - \int_0^{\infty} e^{-\frac{t}{\tau_D(1-E)}} dt\right] = \cdots = D^*_{t=0}\tau_A E \quad \text{(Eq. S26)}$$

As one can examine, the donor fluorescence intensity (eq. S25) decreases monotonically, and the acceptor fluorescence intensity (eq. 26) increases monotonically as the FRET efficiency increases. Additionally, in the presence of donor-only or acceptor-only molecules, many contributions to the overall donor or acceptor fluorescence intensity will be added to the donor or acceptor fluorescence intensities, respectively, solely based on their relative concentration and position relative to the gaussian-shaped PSF. The effect of acceptor-only molecules is not shown in eq. 26, since it describes a pure FRET-based acceptor fluorescence term. Therefore, in a similar fashion to the rationale presented in eq. S23, if the donor-acceptor FRET pair will be at the center of the Gaussian PSF, its contributions to the reduction in donor fluorescence intensity or to the enhancement in acceptor fluorescence intensity will be maximal. However, differences in concentrations within the Gaussian PSF will also influence changes in the overall donor or acceptor fluorescence intensities.

In FRET experiments, it is common to use fluorescence intensity ratios as reporters of FRET. The acceptor fluorescence intensity following donor excitation, normalized to the acceptor fluorescence intensity after direct acceptor excitation (eq. S27).

$$\frac{I_{DA}}{I_{AA}} = \frac{D^*_{t=0}\tau_A E}{A^*_{t=0}\int_0^\infty e^{-\frac{t}{\tau_A}}dt} = \cdots = \frac{D^*_{t=0}}{A^*_{t=0}} \cdot E \quad \text{(Eq. S27)}$$

Therefore, for a pure FRET sample, the ratio of acceptor fluorescence intensity after donor excitation versus after acceptor excitation depends monotonically on the FRET efficiency, and also on other the stoichiometry of donors and acceptors in the sample. Even more so, for a pixel that includes both donor-acceptor FRET pairs and donor- or acceptor-only molecules, this parameter will be influenced by the overall donor and acceptor densities and positions relative to the gaussian-shaped PSF.

Another ratiometric parameter, the ratio of acceptor and donor fluorescence intensities after donor excitation, can be calculated (eq. S28).

$$\frac{I_{DA}}{I_{DD}} = \frac{D^*_{t=0}\tau_A E}{D^*_{t=0}\tau_D(1-E)} = \frac{\tau_A}{\tau_D} \cdot \frac{E}{1-E} \quad \text{(Eq. S28)}$$

As one can see, this ratio increases monotonically with the increase in the FRET efficiency, however without depending on the concentration of donors or acceptors. The lack of dependence of the pure FRET contribution on the donor or acceptor concentrations assists in the interpretation of data acquired for a pixel with both donor-acceptor FRET pairs and donor- and acceptor-only molecules. A base value is expected for non-FRET contributions, and on top of it locally maximal value would suggest maximal contributions to FRET.

<u>Mean photon nanotimes are equivalent to the mean fluorescence lifetime:</u>

Assume the fluorescence decay can be describes as a sum of multiple exponential decays (eq. S29).

$$I(t) = \sum_i \alpha_i e^{-\frac{t}{\tau_i}} \quad \text{(Eq. S29)}$$

where $\alpha_i$ and $\tau_i$ are the amplitudes of these exponential components and their lifetimes, respectively. The intrinsic mean fluorescence lifetime takes the following form (eq. S30).

$$\bar{\tau} = \frac{\sum_i \alpha_i \tau_i^2}{\sum_i \alpha_i \tau_i} \quad \text{(Eq. S30)}$$

It is straightforward to show that the mean photon nanotime of a process that is described with a probability density function of the form of eq. S28 equals the intrinsic mean fluorescence lifetime (eq. S31).

$$\bar{\tau} = \frac{\int_0^\infty \sum_i \tau_i \alpha_i e^{-\frac{t}{\tau_i}} dt}{\int_0^\infty \sum_i \alpha_i e^{-\frac{t}{\tau_i}}} = \frac{\sum_i \alpha_i \int_0^\infty \tau_i e^{-\frac{t}{\tau_i}} dt}{\sum_i \alpha_i \int_0^\infty e^{-\frac{t}{\tau_i}} t} = \frac{\sum_i \alpha_i \left[ (\tau_i^2 - \tau_i t) e^{-\frac{t}{\tau_i}} \right]_0^\infty}{\sum_i \alpha_i \left[ -\tau_i e^{-\frac{t}{\tau_i}} \right]_0^\infty} = \frac{\sum_i \alpha_i \tau_i^2}{\sum_i \alpha_i \tau_i} \quad \text{(Eq. S31)}$$

Importantly, this calculation is ideal since it does not consider nanotimes of background detections, which distribute as if the fluorescence lifetime is infinite. The equivalence of the mean photon nanotime to the intrinsic mean fluorescence lifetime holds as long as the rate of fluorescence photons is much higher than the background rate. In this work, cooled hybrid PMTs, with very low background rate (<100 Hz) was used, and the fluorescence rate is at least 10 KHz, and can be as high as 200 KHz. The fluorescence rate could have been higher than 200 KHz, but this would introduce time-correlated single photon counting pule-up effects, which would distort the underlying fluorescence decays.

### Supplementary Figures

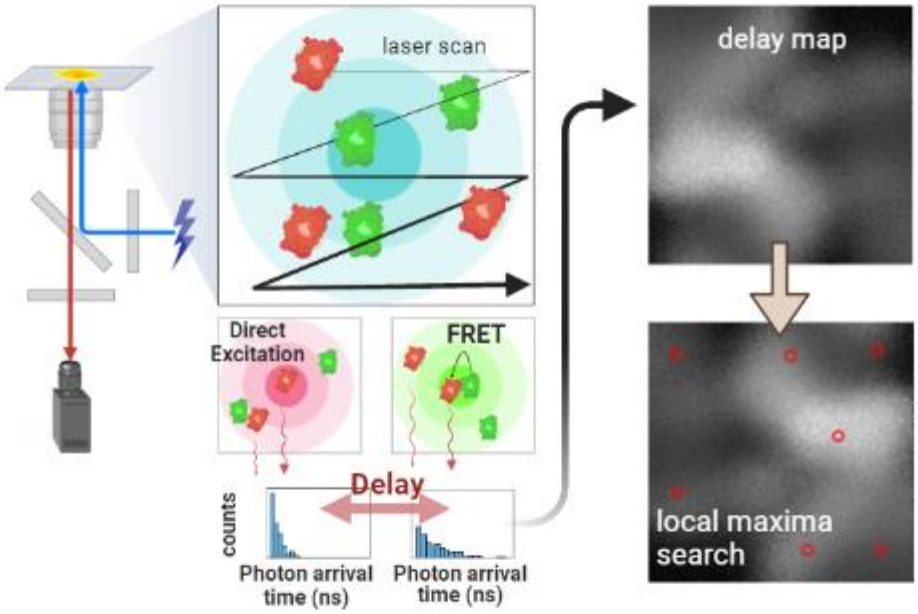


**Fig. S1. The design of simulations to test the FRETsael concept.**

As in a confocal-based FLIM with PIE capabilities, the simulated area is iteratively scanned with a Gaussian-shaped excitation profile. Donor-labeled proteins (green) and acceptor-labeled proteins (red) found in the 2D profile at distances larger than FRET scale will produce acceptor photons only via donor leakage and direct acceptor excitation processes. Alternatively, there are donor-acceptor pairs that are interacting and found within FRET distances, which will produce acceptor photons, detected with a delay. The number of photons produced is dependent on the position of the dye-labeled proteins relative to the PSF profile. By subtracting the images of acceptor mean fluorescence lifetimes after donor excitation (green excitation profile) and after acceptor excitation (red excitation profile) at each pixel, an image of the delay in the acceptor mean fluorescence lifetimes is formed, where pixels with high contribution of FRET interacting pairs will have higher delay values. Next, a local maxima algorithm finds the positions suspected to include the maximal contribution to FRET. A similar design approach is applied also for other measures of FRET, where the position of the maximal contribution to FRET is searched for as a local extremum.

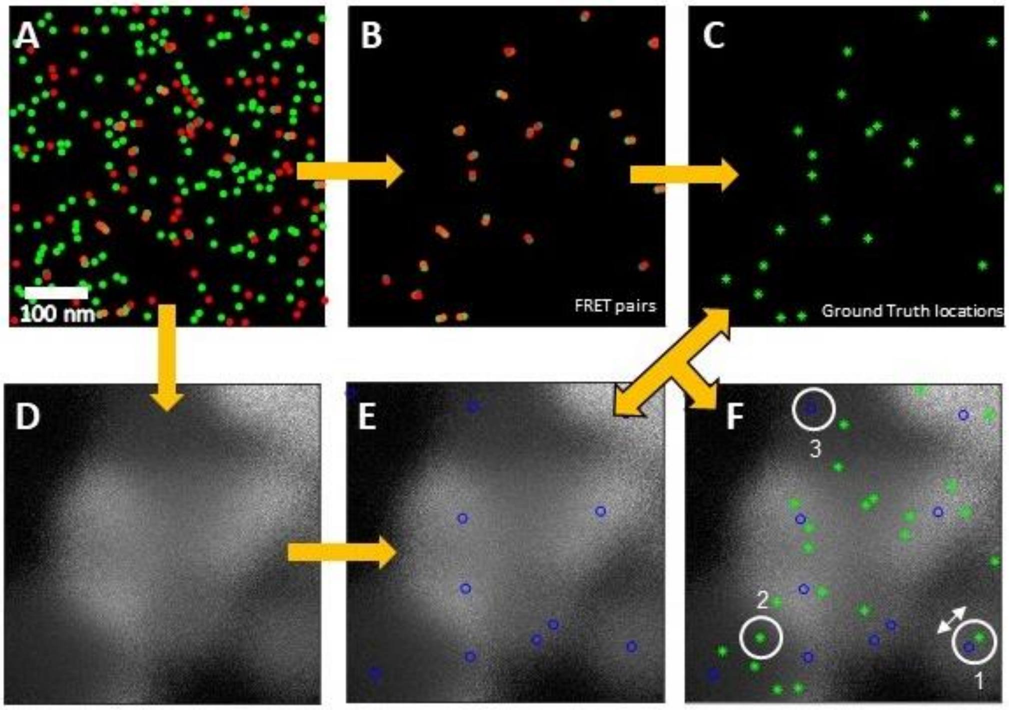


**Fig. S2. The simulation process and its analysis.**

**A**. Donor-labeled proteins (green) and acceptor-labeled proteins (red) are randomly distributed. Orange arrows indicate the pipeline for attaining the ground-truth locations vs. the localizations gathered from FRETsael and the comparison between them. **B**. A fraction of acceptor-labeled proteins are in close proximity to donor-labeled proteins within FRET range (i.e., <1.5 the Förster distance for the given donor-acceptor pair). **C**. The middle position between interacting donor-acceptor pairs is considered as the location that was found for the interaction – a ground-truth location. **D**. The simulation algorithm is iterated on the simulated locations and provides the delay in acceptor mean nanotimes due to FRET for each pixel. **E**. local maxima algorithm locates the positions suspected to be positions for interacting pairs. **F**. By comparing to the ground-truth with the localizations provided by the scanning algorithm the accuracy of the detections is measured (white arrow). Moreover, additional parameters as the number for true positives (circle 1), false negative (circle 2) and false positive (circle 3) events is provided.

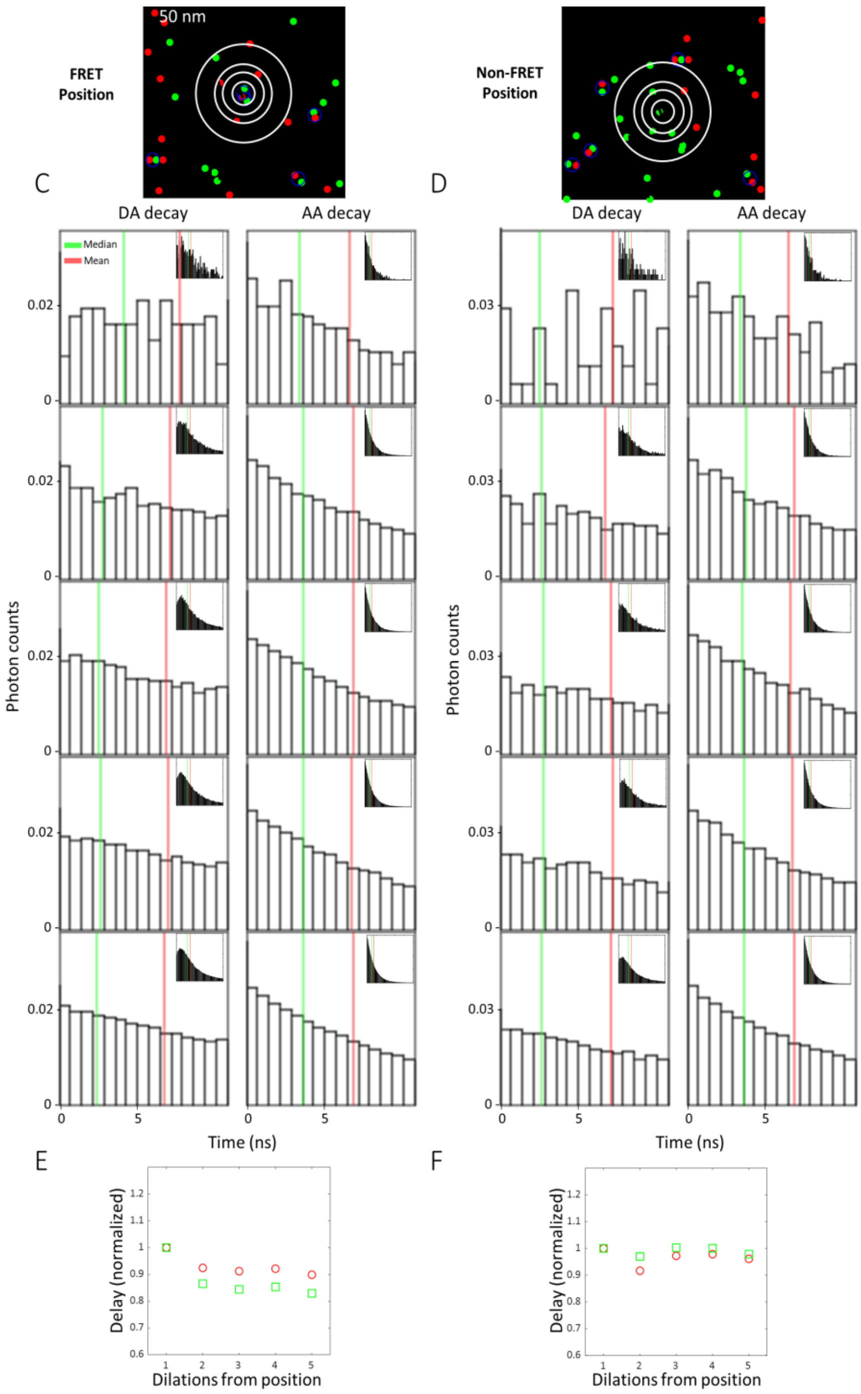


**Fig. S3. Simulations of photon nanotime histograms at different dilations away from FRET and non-FRET positions.**

**A, B**. Donor-acceptor FRET pair in the simulations (**A**) and donor only (**B**) positions, surrounded by circles of dilations from positions. **C, D**. Zoomed-in photon nanotime histograms in DA channel (donor excitation acceptor emission; left) and in AA channel (acceptor excitation acceptor emission; right) for pixels located over the circles depicted in A or B. Inset shows the entire range of the photon nanotime histograms. In the case of FRET (**C**), or non-FRET donor-only (**D**), green and red vertical lines highlight the median and mean of photon nanotimes, respectively. **E, F**. The median of all photon nanotimes (green boxes) and the mean fluorescence lifetimes (red circles) as a function of the dilations from the positions of the FRET pair (**E**) or of the non-FRET donor-only (**F**).

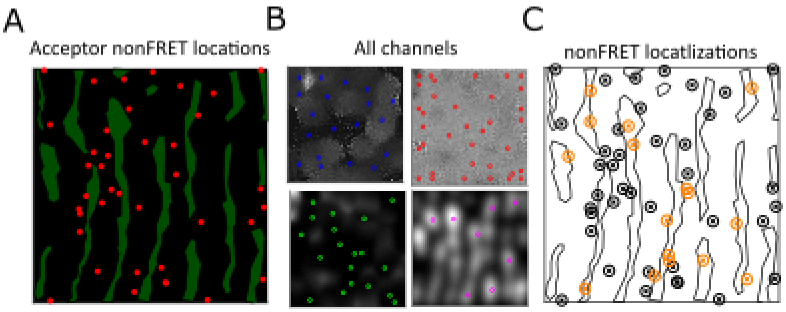


**Fig. S4. ER-Ribosomes simulation for non-FRET localizations.**

**A**. Red dots (#80) representing ribosomes that were spread far from the ER green patches **B**. The output of the four parameter channels with localizations in each parameter channel (delay in the acceptor mean fluorescence lifetime in blue; donor mean fluorescence lifetime in red; acceptor to donor fluorescence intensity ratio in green; donor fluorescence intensity in pink). **C**. The ground-truth of non-interacting ribosomes represented as black ⊗ signs. Localizations of suspected areas with no interaction (orange ⊗ signed) presented next to the ground-truth locations. Comparing these localizations with true locations yielded detection accuracy of 67±32 nm (SD) with TPR of 18% and the FDR of 56% (CSI=0.14).

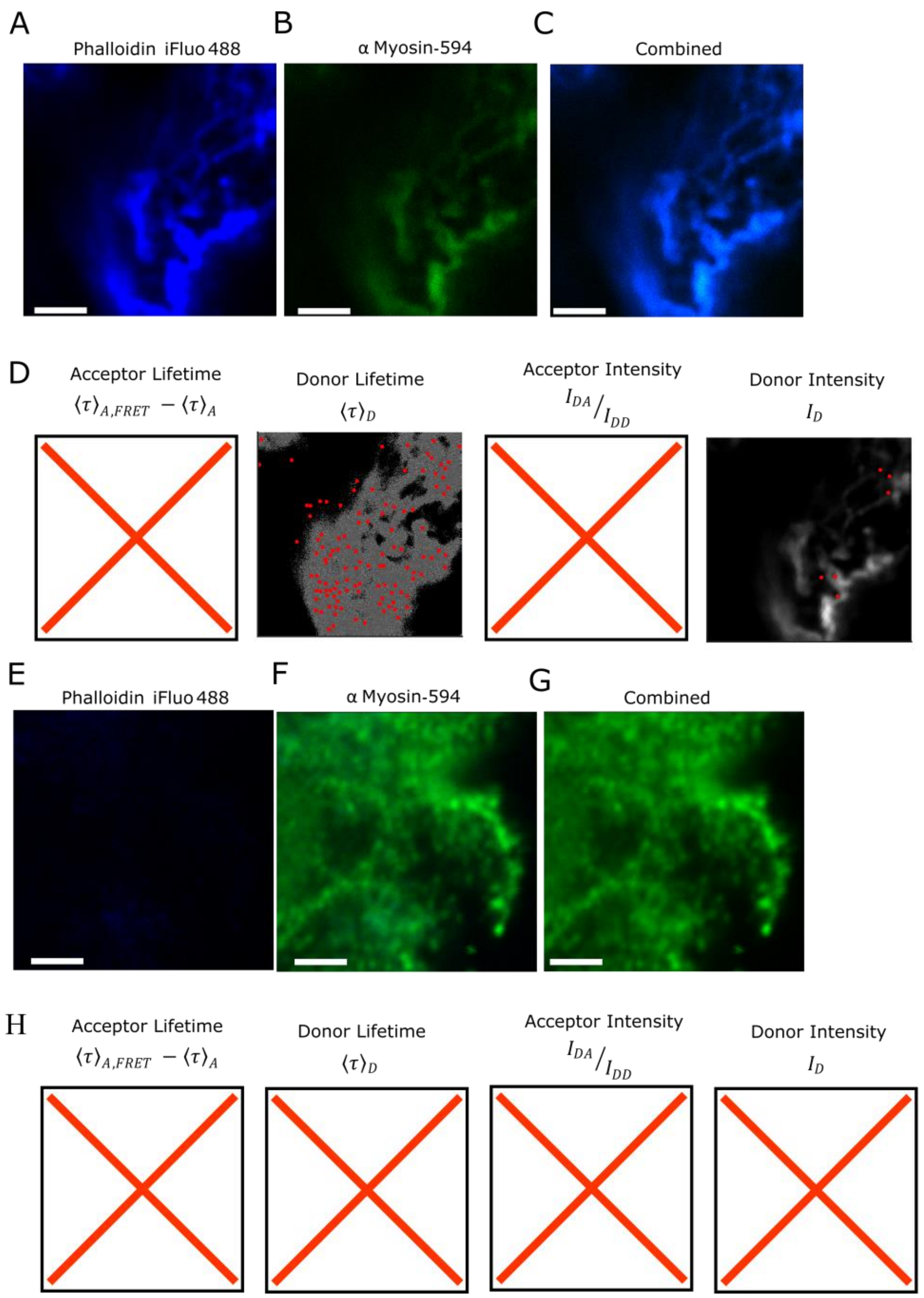


**Fig. S5. Controls for FRET imaging of donor-only iFluor 488-phalloidin and acceptor-only Alexa Fluor 594-anti-Myosin antibodies in SH-SY5Y cells.**

**A, B**. Blue and green fluorescence detection channels for fluorescence intensity images of iFluor 488 conjugated to phalloidin, shown in two detection channels. **C**. Merge of panels **A** and **B**. 10x10 μm$^2$, 512x512 pixels. Scale bar 1 μm. **D**. Only two of the four different parameter channels yielded localizations (donor mean fluorescence lifetime; donor fluorescence intensity). Localizations are depicted in red circles. **E, F**. Blue and green fluorescence detection channels for fluorescence intensity images of Alexa Fluor 594 conjugated fluorescent antibodies binding non-muscle myosin IIA, shown in two detection channels. **G**. Merge of panels **E** and **F**. 10x10 μm$^2$, 512x512 pixels. Scale bar 1 μm. **H**. None of the parameter channels yielded localizations. Panels with X signs signify parameter channels in which the FRETsael algorithm did not identify localizations.

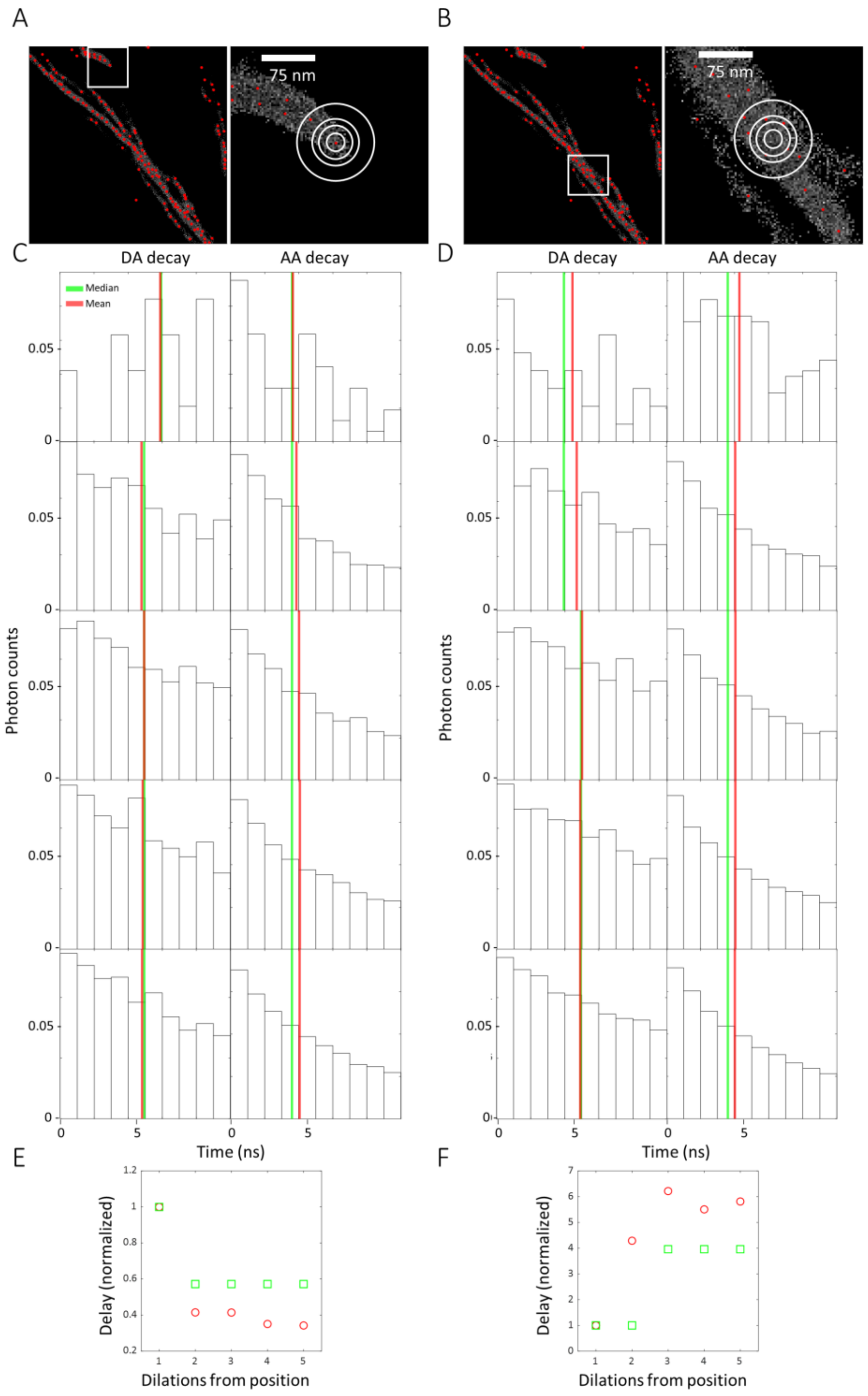


**Fig. S6. Experimental photon nanotime histograms.**

**A, B**. A localization extrapolated from the FRETsael algorithm (**A**) and a random location inside the cell (**B**), surrounded by circles of dilations from the localization positions. **C, D**. Photon nanotime histograms in DA channel (donor excitation acceptor emission; left) and in AA channel (acceptor excitation acceptor emission; right) for pixels located over the circles depicted in A and B. In the case of localization (**C**) or random location (**D**) green and red vertical lines highlight the median and mean of photon nanotimes, respectively. **E, F**. The median of all photon nanotimes (green boxes) and the mean fluorescence lifetimes (red circles) as a function of the dilations from the positions of the localization (FRET pair) (**E**), or a random location case (**F**).
